# Engineered sensor bacteria evolve master-level gameplay through accelerated adaptation

**DOI:** 10.1101/2022.04.22.489191

**Authors:** Satya Prakash, Adrian Racovita, Clenira Varela, Mark Walsh, Roberto Galizi, Mark Isalan, Alfonso Jaramillo

## Abstract

Gene circuits enable cells to make decisions by controlling the expression of genes in reaction to specific environmental factors^1^. These circuits can be designed to encode logical operations^2–7^, but implementation of more complex algorithms has proved more challenging. Directed evolution optimizes gene circuits^8^ without the need for design knowledge^9^, but adjusting multiple genes and conditions^10^ in genotype searches poses challenges^11^. Here we show a multicellular sensor system, AdaptoCells, in Escherichia coli, that can evolve complex behavior through an accelerated adaptation to chemical environments. AdaptoCells recognize chemical patterns and act as a decision-making system. Using an iterative improvement method, we demonstrate that the AdaptoCells can evolve to achieve mastery in the game of tic-tac-toe, demonstrating an unprecedented level of complexity for engineered living cells. We provide an effective and straightforward way to encode complexity in gene circuits, allowing for fast adaptation in response to dynamic environments and leading to optimal decisions.

---

Gene circuits are a fundamental tool in biotechnology, allowing regulation of gene expression and enabling organisms to respond to environmental changes using chemical sensors. However, despite progress in gene circuit design^12^, it remains a challenge to create circuits that can make complex decisions, such as playing an expert strategy in a board game. This requires a decision-making algorithm based on environmental conditions, represented by chemical sensors and a truth table that determines the optimal move. Conventional gene circuits struggle to encode the truth table of even the simplest board games, due to the need for numerous logic gates and transcription factors.

A promising approach to engineering complex gene circuits is the use of multicellular gene circuits^3, 4^, which allow for communication between cells. This approach involves dividing genes into separate cells within a population, resulting in the collective behavior of the population resembling that of a single-cell gene circuit. By using this divide-and-conquer strategy, it is possible to design more complex gene circuits that mimic internal gene-gene regulations through the engineering of orthogonal cell-cell communications. However, despite this potential, implementing gene-gene regulations at the multicellular level remains a challenging problem due to the limited number of independent chemical communication channels available and the complexity involved in connecting multiple logic gates through internal gene-gene interactions. So far, it has only been achieved for a limited number of independent chemical communication channels^13^, which is not enough to support the design of advanced decision-making that requires multiple logic gates to be connected through internal gene-gene interactions. This limitation hinders the assembly of the numerous logic gates needed for complex truth tables. For example, an implementation of an expert tic-tac-toe player using DNA computing and logic gates required 128 three-input gates^14^.

Analog gene circuits^15, 16^ use continuous signals to perform complex algorithms, monitoring the environment, regulating gene expression, and processing signals. To make these circuits more versatile and enable decision-making, discretizing output genes transforms the continuous output into a discrete or digital signal by dividing it into distinct levels. Although analog gene circuits with discretized output can be useful, gene circuits with digital behavior are more commonly used. The "winner-take-all" strategy is a powerful discretization method that enables decision-making based on the gene circuit with the strongest phenotype^17^. This approach provides several advantages over other discretization methods, as it considers the relative strengths of multiple signals to decide, rather than relying on a single threshold value. However, designing analog gene circuits for decision-making can still present challenges due to their sensitivity to gene expression levels of each transcription factor. Automated design methods, such as computational design, have aided the engineering of analog gene circuits for biological adaptive behaviors^18^, but they have not yet successfully implemented advanced decision-making^19, 20^. Nonetheless, this has been achieved artificially using physical and chemical systems by engineering memory units with neuromorphic computing capabilities^21–27^.

Darwinian directed evolution’s trial-and-error approach - that generates genetic variations like mutations or recombination - has proven to be a challenge for engineering complex gene circuits in living cells. The lack of continuity in phenotype space, slow pace, and difficulty in covering the vast genetic space of multi-gene circuits limit its effectiveness. Furthermore, the multi-objective selection process for truth tables in gene circuits implementing logic gates often involves complex methods and high concentrations of antibiotics, reducing genetic diversity. The challenges of discontinuity in phenotype space, slow pace, and difficulty in covering the vast genetic space of multi-gene circuits hinder the use of Darwinian directed evolution for engineering advanced decision-making capabilities in living cells.

Here we report a new strategy for engineering complex multicellular gene circuits in living cells by developing a cellular adaptive sensor system, AdaptoCell, using modified *E. coli* cells. Our approach incorporates gene regulation into a synthetic plasmid heteroplasmy^28^ system that allows cells to maintain and adjust the expression of a red fluorescent protein and a specific resistance protein across generations. We engineer nine AdaptoCells, each sensing a different chemical. When grown together, they adjust their red fluorescence expression to match a specific nine-chemical pattern. We link the expression of the red fluorescent protein to a kanamycin resistance protein so that as the resistance protein increases, the red fluorescence decreases. This approach enables the use of sub-lethal kanamycin concentrations to permanently adapt the cells, resulting in lower expression levels of the red fluorescent protein. By choosing the AdaptoCells exhibiting the highest fluorescence, we can implement a decision-making process. This process involves exposing the cells to various chemical patterns, and if the decision produces an undesired outcome, we induce adaptation in the cells responsible for the incorrect decision. This iterative improvement method is repeated until the desired decision is achieved. We demonstrate this method by applying it to simple board games such as tic-tac-toe, a common benchmark in artificial intelligence^29^. We show that AdaptoCells can evolve to a state where they never lose a game. In summary, we report the creation and evolution of multicellular sensor systems called AdaptoCells, which can recognize and make decisions based on chemical patterns, as demonstrated through the example of mastering board games using an iterative improvement method.

## Results

### Sensor cells with synthetic plasmid heteroplasmy

We have advanced the concept of synthetic plasmid heteroplasmy by incorporating a conditional mechanism based on an external inducer. Synthetic plasmid heteroplasmy, as initially proposed^28^, enables the stable maintenance of two plasmids in a population of *E. coli* cells with the ability to modify their relative copy numbers through encoded genetic machinery. This system is based on engineering multi-copy plasmids that carry the same antibiotic resistance gene, analogous to heterozygotic mutations observed in multi-copy plasmids^30^ or mitochondrial DNA^31^.To maintain plasmid stability, both plasmids must have the same replication rate and genetic load, as differences can result in the loss of one plasmid type. However, maintaining plasmid composition across generations can be challenging when both plasmids express different gene expression cassettes in a constitutive manner^28^, which prevents self-dosage. To overcome this limitation, we have improved synthetic plasmid heteroplasmy by making it conditional on an external inducer. This allows gene expression from both plasmids to switch off during normal cell growth and activate only when desired, resulting in equal genetic loads during growth and enhanced plasmid stability. Additionally, this conditional mechanism facilitates adaptation to chemical environments by allowing cells to alter their plasmid content after sensing a specific chemical inducer.

To engineer our synthetic heteroplasmy, we utilized two multicopy plasmids, P1 and P2, in populations of *Escherichia coli* Marionette strains (Fig. 1a). Both plasmids carried the same ampicillin resistance gene (AmpR) and origin of replication, regulating the stability of the total plasmid copy number (*a+b*), but not the individual copy numbers of P1 and P2. To ensure the stability of the individual copy numbers of P1 (*a*) and P2 (*b*), we designed the plasmids to have the same length and genetic load. We referred to the relative frequency of P1 in the cell as "weight", which was calculated as *a/(a+b)*. This weight serves as a genotype, as it maintains stability and consistency across generations, enabling us to predict the transmission of population-level traits and characteristics from parent to offspring. The plasmids encode a promoter from the Marionette cassette, enabling us to sense chemicals by using inducible promoters without any significant cross-interference, to regulate the genetic cargo of the plasmid heteroplasmy.

**Fig. 1.**
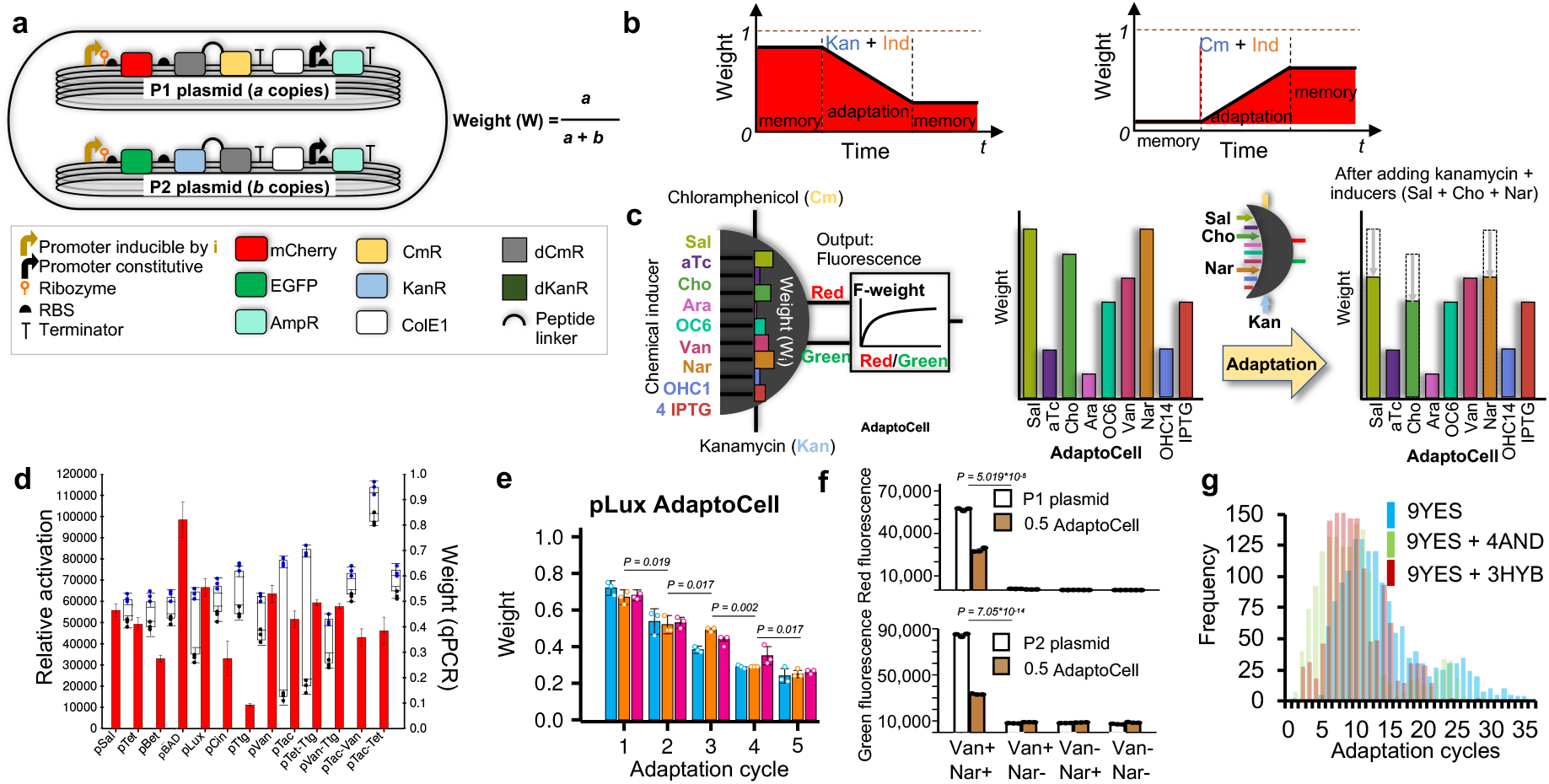
Engineering stable genotypes in Escherichia coli through synthetic plasmid heteroplasmy. **a**, We combined two multicopy plasmids, P1 and P2, to create a synthetic plasmid heteroplasmy. P1 and P2 carry an ampicillin resistance marker (AmpR) and a common replicon (ColE1) regulating the stability of the total copy number (a+b), but not the individual copy numbers of P1 and P2 (a and b). In addition, the stability of a and b, is maintained by designing the plasmids to produce equal genetic loads during replication preserving the relative frequency of P1 in the cell, referred to as "weight". By incorporating mCherry into P1 and EGFP into P2, we can determine the weight through the fluorescence ratio of the proteins when the promoters are fully active. To equip the cells with the ability to detect chemicals, we integrated the same inducible Marionette promoter into both plasmids. This promoter, triggered by the cognate chemical inducer "i," regulates the operon containing fluorescent protein and antibiotic resistance protein fusions (KanR/CmR for kanamycin/chloramphenicol resistance) or their non-functional counterparts (dKanR/dCmR). We implement a red fluorescence phenotype in AdaptoCells through engineering a genetic link between the expression of a kanamycin resistance protein and a red fluorescent protein. As the expression of the kanamycin protein increases, the red fluorescence decreases. **b**, The selection process using Kanamycin (Kan) or Chloramphenicol (Cm) amplifies the fraction of cells carrying the P2 or P1 plasmid (coding for KanR or CmR), which respectively reduces or increases the cell population’s weight, but only when KanR or CmR are induced with a cognate chemical (ind). **c**, Antibiotic adaptation in AdaptoCells enables the simultaneous adjustment of multiple weights when adding the antibiotic in an environment representing a chemical pattern. For instance, the diagram illustrates the kanamycin adaptation of AdaptoCells when exposed to a chemical environment featuring high levels of Sal, Cho, and Nar inducers. The kanamycin adaptation only reduces the weight of the induced strains, leading to a lower red fluorescence output and F-weight when encountering again environments with the same Sal, Cho, and Nar levels. **d**, Characterisation in fluorescence and weight of the 9YES and 4AND AdaptoCell libraries containing 9 single-input and 4 dual-input promoters, respectively. Fluorescence variation upon induction (relative activation) of each AdaptoCell (red bars, left y-axis). Weight shown (right y-axis) before (blue) and after (black) one cycle of adaptation with kanamycin. **e**, Example of a 9YES AdaptoCell (pLux) of 0.6 weight, characterised after 5 consecutive adaptation cycles (see Extended Data Fig. 3 and 4 for the equivalent plots from flow cytometry, DNA sequencing and qPCR, for all the AdaptoCells). **f,** Normalised red and green fluorescence of one member of the 4AND AdaptoCell library (constructed by re-engineering the pVan promoter with the TtgR operator; see Methods and Extended Data Fig. 2b for the 3 other 4AND re-engineered promoters), with 0.5 weight, in the presence and absence of Vanillic acid (Van) and Naringenin (Nar) inducers. **g**, Distribution of the kanamycin adaptation cycles required to achieve mastery (100% expertise) in 1,500 3×3 board games for the 9YES, 9YES + 4AND and 3HYB (containing 3 hybrid AdaptoCells, see Box 1) AdaptoCell libraries. (data show mean ± s.d. for n=3 independent biological replicates).

To easily measure the weight, we incorporated mCherry into the cargoes of P1 and EGFP into P2, enabling us to determine the weight through the fluorescence ratio of the mCherry and EGFP proteins. Additionally, we equipped the plasmids P1 and P2 with the ability to activate the fluorescence proteins and specific antibiotic resistance proteins after sensing a specific chemical inducer (Fig. 1a). Both plasmids encode the same inducible promoter, which regulates an operon with the fluorescent proteins and the fusions of antibiotic resistance proteins (KanR/CmR for kanamycin/chloramphenicol resistance) or their respective non-functional "dead" forms (dKanR/dCmR). We determine the weights through fluorescence on a plate reader, flow cytometry, DNA quantification with sequencing^32^, or qPCR. Our results confirm that the population’s weight stays consistent for several days (see Extended Data Figs. 1a-c, 3b-d and Supplementary Text), aligning with a prior report^28^.

### Sub-lethal antibiotic selection produces adaptation in plasmid heteroplasmy

We hypothesized that the introduction of sub-lethal antibiotic concentrations would result in a biased alteration of the distribution, as cells harboring a higher number of plasmids encoding an antibiotic resistance protein will become enriched (in conditions where the appropriate inducible promoter is also chemically induced to express the antibiotic resistance gene). For instance, the combination of kanamycin and the chemical inducer allowed us to select cells that had the highest number of P2 plasmids. This is because the P2 plasmids carry the gene that makes the cells resistant to kanamycin and this gene is controlled by an inducible promoter. As a result of the selection process, the average number of P2 plasmids increased while the average number of P1 plasmids decreased. However, the total number of plasmids (P1 and P2 combined) in the cells remained constant. This shift towards cells with fewer P1 plasmids caused a decrease in the average red fluorescence per cell. The P1 plasmid is responsible for producing the red fluorescence protein mCherry. The simultaneous addition of both the chemical inducer and antibiotic triggers the cells to alter their weight, which in turn changes their fluorescence and antibiotic sensitivity activities. By having the P1 and P2 plasmids contain a different inducible antibiotic resistance gene, the addition of an antibiotic shifts the plasmid ratio towards the plasmid with the appropriate resistance gene. Cells with higher numbers of P1 plasmid copies exhibit higher levels of red fluorescence when the promoter is induced, but they are also more sensitive to kanamycin. This is because they possess an inactive kanamycin resistance gene (dKanR), while the active kanamycin resistance gene (KanR) resides on the P2 plasmid. The presence of kanamycin and the chemical inducer prompts these cells to decrease their population averaged P1 copy number, leading to a reduction in red fluorescence levels. The total red fluorescence of AdaptoCells reflects the weighted average fluorescence among induced strains. When kanamycin is added to an environment containing several of the nine chemical inducers, it lowers the weights of the induced strains, causing a decrease in total red fluorescence in that environment, but not in others (Fig. 1c). Thus, the total red fluorescence can be modulated among different environments by trial-and-error only through kanamycin adaptation, which lowers the red fluorescence in the targeted environments.

An AdaptoCell constitutes in fact a minimal gene circuit able to adapt its gene expression when it is active, analogous to the memristor element used in electronic circuits with neural network behaviour^33^. The promoter and KanR act as the computational and adaptation engines respectively, where the promoter could be replaced by a more complex genetic logic gate (Fig. 1d) (see Box 1 for generalisations).

### AdaptoCells libraries allow for the adaptation to chemical patterns

We chose 9 different Marionette promoters to regulate the AdaptoCell’s fluorescent proteins, thus creating a base AdaptoCell library of 9 single-input AdaptoCells (9YES library, characterised in Fig. 1d). We also extended the base library by re-engineering 4 Marionette single-input promoters to have an additional input. These conditionally transcribe when two specific chemicals are present in appropriate concentrations, implementing an AND logic gate. We call this library 4AND (see Fig. 1d,f and Extended Data Fig. 2b for their characterisation). After induction with the cognate chemical(s), all the 9YES and 4AND AdaptoCells showed a significant activation versus non-induced (or partially induced) states (Fig. 1d, average ANOSIM of R=0.82, see Suppl. Table S5). To experimentally test a generalisation of AdaptoCells (Box 1) for constructing other adaptive gene circuits, we built a hybrid AdaptoCell (pBAD:pSal) whose P1 and P2 plasmids encode for the AraC-mediated activation and LysR-mediated repression of the pBAD and pSal promoters respectively. Its functionality interpolates between a YES gate (0 and 1 weights) and an OR logic gate of the promoters. We also constructed pTet:pBAD and pTet:pVan, creating the 3HYB library (Extended Data Fig. 2c).

**Fig. 2.**
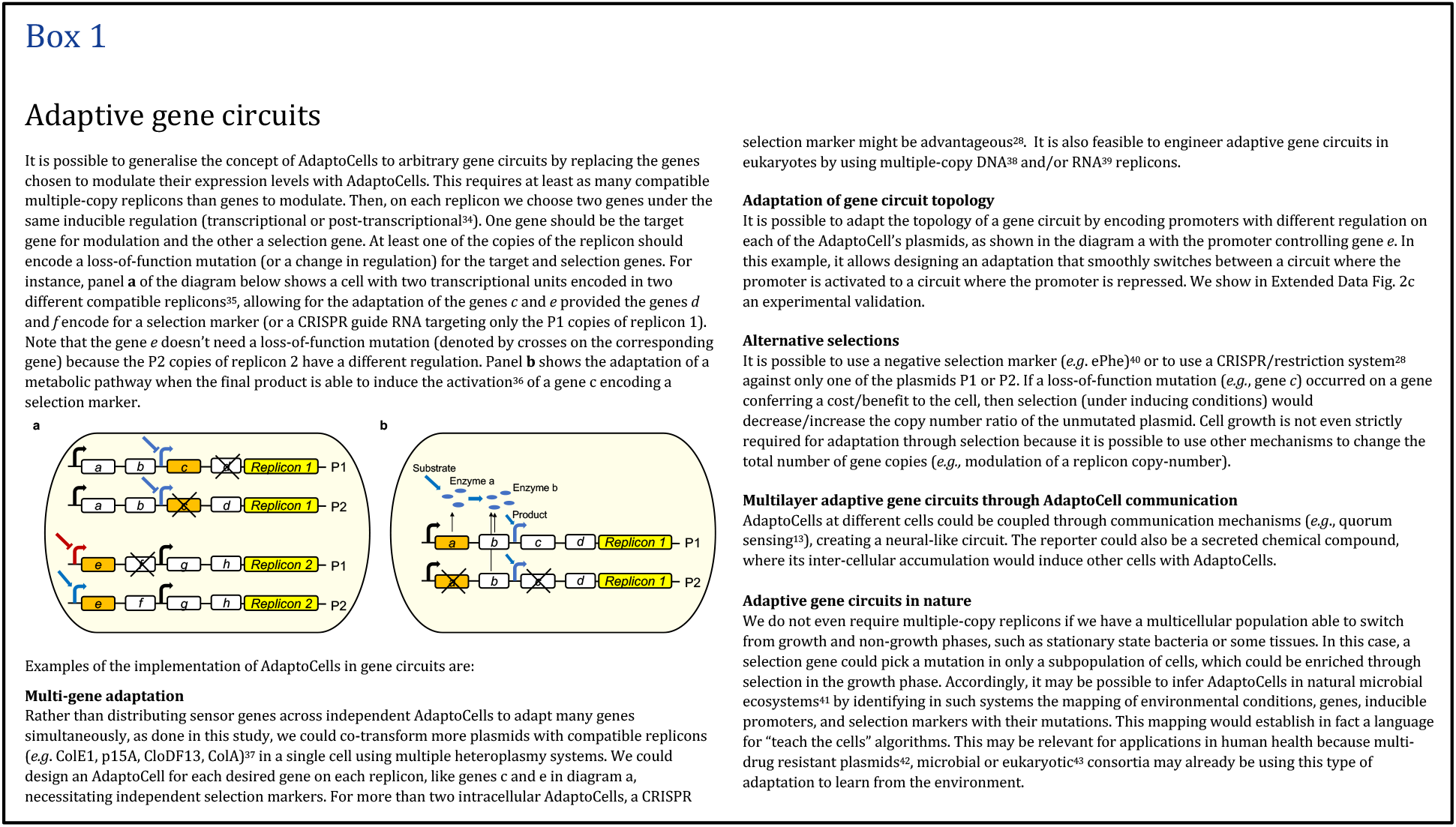
**Box 1.**

To analyse the capability of our libraries to successively adapt throughout several cycles under the cognate inducer, we exposed our 9YES and 4AND libraries to consecutive cycles of adaptation (Extended Data Fig. 3a). Each adaptation cycle involved overnight growth from frozen stock, growth in presence of kanamycin and the respective inducers, and finally the creation of another glycerol stock. After each cycle, the weights decreased significantly (p<0.05), indicating adaptation. This was further validated by analysing the single-cell weight distributions using flow cytometry (Extended Data Fig. 5a), where we inferred the weight from a ratiometric analysis of the green and red fluorescence values (Methods and Supplementary Text). The mean values of the distributions quantitatively agree with the results obtained through plate reader fluorescence assay (average R^2^ 0.91, Extended Data Fig. 4c). Our weight values after adaptation correlated with a model prediction of the effect of adaptation on the single-cell weight distributions (R^2^ 0.92, Extended Data Fig. 5b). The model only assumed that the antibiotic killed the cells having a red fluorescence higher than a given threshold (Supplementary Text).

**Fig. 3.**
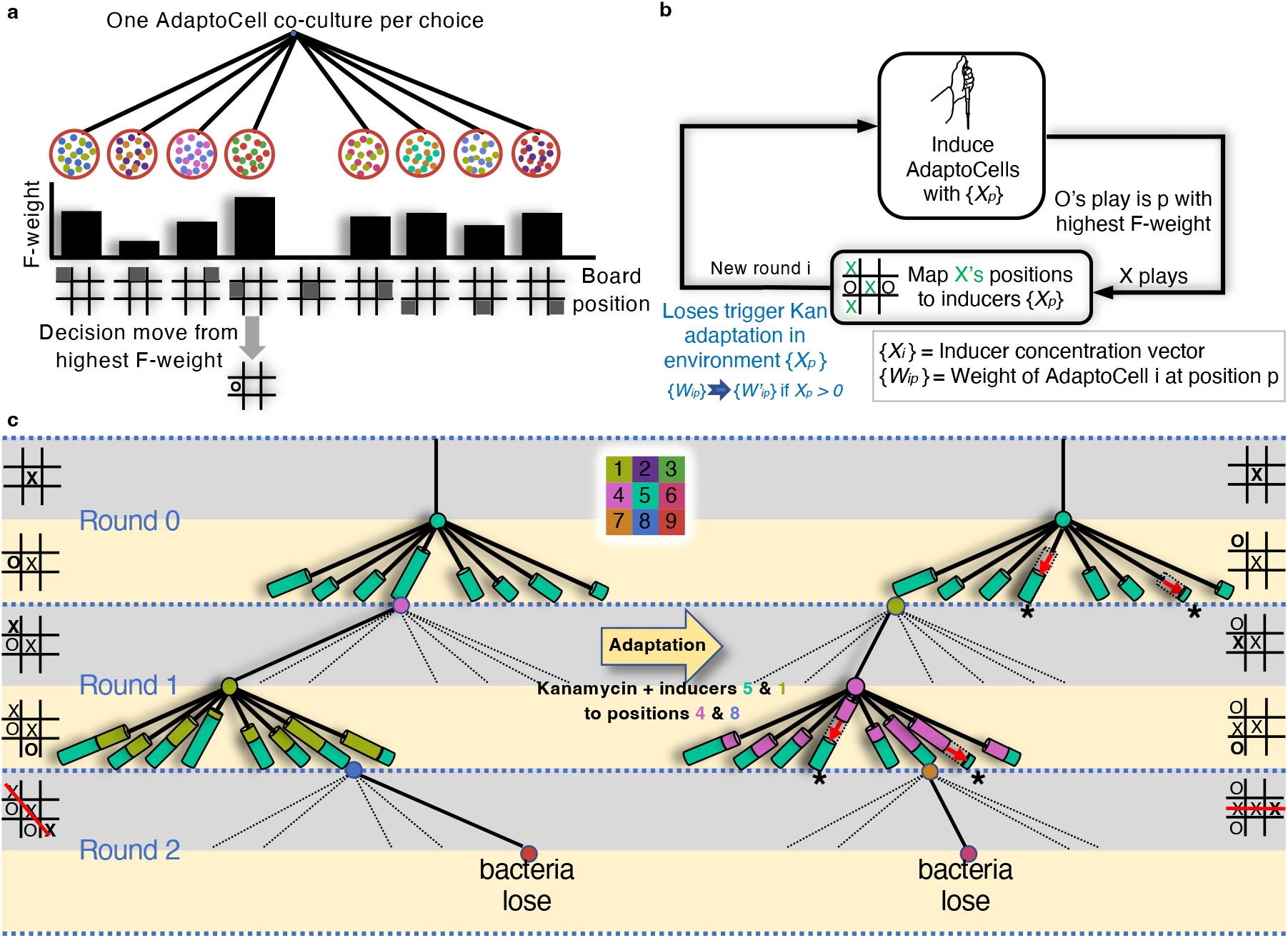
Decisions in tic-tac-toe game from chemical Induction and adaptation. **a**, The outcome of each round in a tic-tac-toe game is determined by incubating the AdaptoCells co-cultures with the inducers associated with the X player’s moves, and measuring the highest F-weight to choose the bacteria’s move. **b**, Diagram showing the iterative adaptation process that begins with growing the AdaptoCells in an environment defined by the inducers mapped to the opponent’s moves. The environment changes with each round as the opponent makes their move, which makes the cells respond differently. Only when player O loses, we perform a kanamycin adaptation in presence of the environment created during the losing match, which changes the state of the cells from O_i_ to O_i+1_. **c**, Decision tree for the tic-tac-toe game shown as branches with cylinders of varying height to represent the weights for each strain of bacteria associated with a specific inducer, which represents the X player’s move. In round 0, the branches show the weights for strain 5, which corresponds to inducer 5 (X player’s position), at all possible positions. After inducing the co-cultures with chemical 5, the highest F-weight is the branch with the tallest cylinder height, representing the move of the bacterial player O. The X player then choses to move to position 1. In the following round, the cocultures at unoccupied positions are induced with all the inducers corresponding to X’s positions, which in this case are chemicals 5 and 1. This creates an environment with inducers 5 and 1, so the weights for strains 5 and 1 are represented as stacked cylinders in the branches associated with all unoccupied positions for the bacterial player O. The game ends in the next round when X plays 9 and wins. Afterwards, a kanamycin selection operation is applied to only the cocultures at the positions played by player O (positions 4 and 8) in the presence of the inducers of X’s moves from all rounds before the win (inducers 5 and 1). The X player’s last move is not considered in this process because it did not create an environment to grow the AdaptoCells bacteria.

**Fig. 4.**
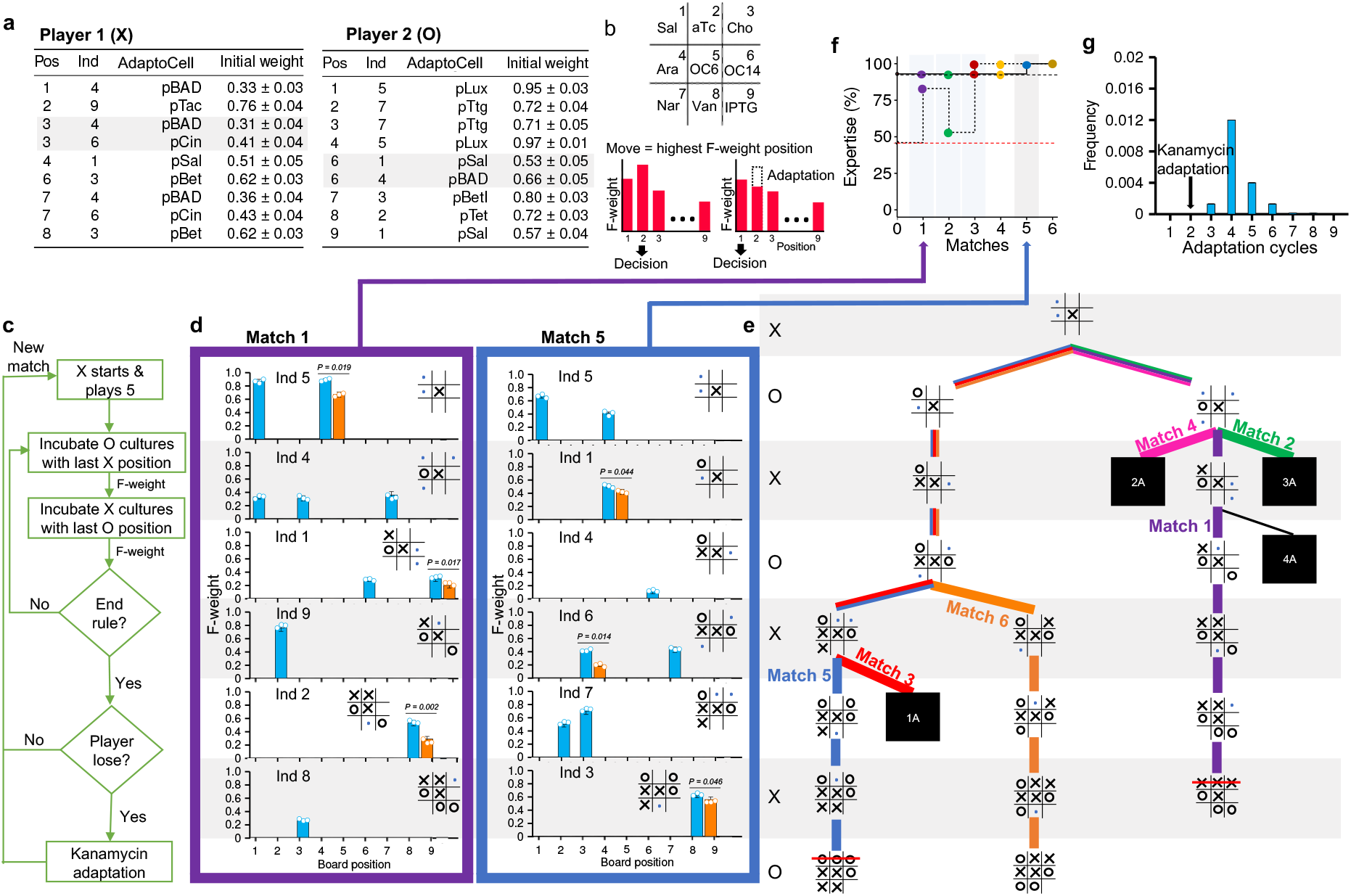
Two bacterial libraries undergo adaptation to master Tic-Tac-Toe by playing against one another. **a**. Libraries of AdaptoCell *E. coli* cultures designed to implement a restricted decision tree. We show the board position (Pos) and the number of the chemical (Ind) activating the AdaptoCell’s promoter. The last column shows the weights measured by DNA sequencing before adaptation. Positions 3 and 7 for Player 1 and position 6 for Player 2 are assigned multi-cellular co-cultures of two AdaptoCells (Methods). **b,** Chemical inducers are mapped to board positions. **c**, Experimental workflow used in the matches. **d**, Two matches where kanamycin adaptation occurs, showing the board state before making a move. F-weights from AdaptoCell cultures before (blue) and after (orange) kanamycin adaptation (± s.d. for n = 3). Game board insets indicate the last move of the opponent and the permissible positions (blue dots) for the current move, restricted to limit possibilities. We do not allow for all unoccupied positions to restrict the number of possibilities, so we chose a few permissible positions. The position with highest F-weight gives the current move (shown aligned in panel **e**). **e**. The simplified tic-tac-toe decision tree, black boxes represent collapsed branches (Supplementary Text). Horizontal grey overlays align the decision tree to the decision taken by the bacteria in panel **d**. **f**, Expertise acquired during a tournament of 6 matches with both players suffering kanamycin adaptation. Filled circles denote the expertise (main text) after each game or after adaptation, colour-matching the frames in panel **d** (dotted line = player 2). Coloured overlays indicate the learner (grey = Player 1; blue = Player 2). Dashed lines indicate the expertise of non-learners (black = Player 1; red = Player 2). **g**, Histogram of the adaptation rounds required to achieve mastery in a computer simulation of 10,000 random tournaments, where for adaptation we used randomised inducers and kanamycin.

**Fig. 5.**
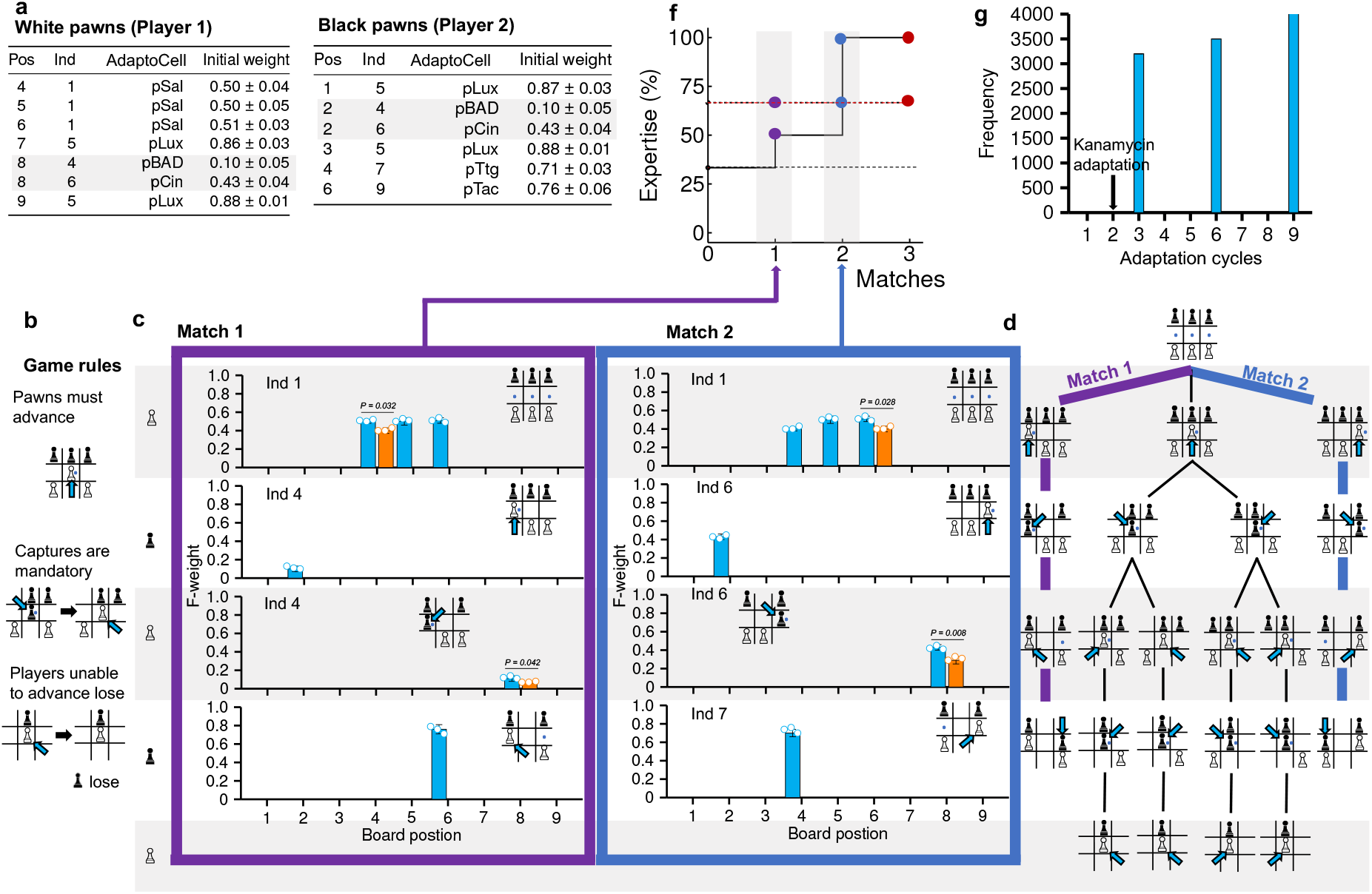
Two bacterial libraries playing and adaptation a simplified chess game in a 3×3 board (hexapawn). **a,** Libraries of AdaptoCell *E. coli* cultures chosen for the decision tree. F-weights were measured before adaptation. Position 8 for Player 1 and position 2 for Player 2 are assigned co-cultures of two AdaptoCells (Methods). **b,** Rules of the game. **c**, Losses of the white pawns lead to kanamycin adaptation with kanamycin (F-weight before (blue) and after (orange) kanamycin adaptation (± s.d. for n = 3)). **d,** Decision tree for the AdaptoCell libraries. Blue dots indicate the available/mandatory moves. **e,** Expertise acquired during the tournaments with kanamycin adaptation after losing a game, (dotted line = player 2). Grey overlays indicate the adaptations of Player 1. Dashed lines indicate the expertise of non-learners (black = Player 1; red = Player 2). **f**, Distribution of adaptation rounds required to achieve mastery for a set of 10,000 simulations of tournaments where the adaptation used randomised inducers and kanamycin (similarly to Fig. 1g).

Now that we had libraries able undergo several rounds of adaptation, we asked to which extent we could predict mCherry expression from an AdaptoCell’s weight. We found that the red fluorescence for 50 AdaptoCells of different weights from the 9YES and 4AND systems, correlated well with weight obtained from DNA quantification (R^2^ 0.93, Extended Data Fig. 3a). We therefore developed a computational platform to simulate the decisions made by our AdaptoCell libraries when subjected to successive adaptation steps. This allowed us to examine if we could use our 9YES AdaptoCell libraries to construct a library of cells able to learn any game played on a 3×3 board in a few adaptations. Given a set of weights, expertise is defined as the percentage of positive results (wins and draws) assessed computationally against all possible games. When we used a 9YES library for each position (all AdaptoCells having equal weights, which implements a player with complete random decisions) to play 1,500 different 3×3 board games, with kanamycin adaptation, we got mastery (100% expertise) in less than 33 adaptations and often in a handful of adaptations (Fig. 1g, Extended Data Fig. 10a,b, and Supplementary Text). Interestingly, when we combined the 9YES and 4AND libraries, we reduced the mean of the adaptation rounds required to achieve expertise from 11 to 6 and the maximum from 33 to 25. When we combined the 9YES and 3HYB libraries, we reduced the mean of adaptation rounds from 11 to 8 and the maximum from 33 to 27 (Fig. 1g). Hybrid AdaptoCells also allow increasing the computation capabilities without relying on the difficult engineering of combinatorial promoters.

### Antibiotic adaptation to change AdaptoCells decisions

By using multiple AdaptoCells, we can create a decision-making system. We define a decision among N categories by choosing the AdaptoCell with the highest red fluorescence among N independent AdaptoCells populations. For instance, in a 3×3 board game, we select the highest F-weight from the AdaptoCell populations at each available board position (Fig. 3a). To optimize the behavior of this system, we have designed an iterative improvement method. For example, when deciding among patterns of occupied board positions, we start by growing an initial set of N AdaptoCells independently. We then interrogate the system’s decision by growing the N AdaptoCells in presence of the equivalent chemical pattern, either with saturating or vanishing levels of chemicals (Fig. 3b, where N=8). If the decision is not the desired one, we perform a kanamycin adaptation only to the AdaptoCells responsible for the decision (i.e., producing the highest red fluorescence). This process generates a new set of N AdaptoCells with updated AdaptoCells that underwent adaptation. We continue this process of adaptation until the decision changes. The adaptation always lowers the red fluorescence, and since we consider the highest red fluorescence to decide, the decision will eventually change.

By strategically targeting the kanamycin adaptation to only the population with the highest F-weight, we cause a change in the decision, as the chosen culture’s F-weight decreases and is no longer the highest. When a decision is not right in a given environment, we perform kanamycin adaptation with the AdaptoCells that have the highest F-weight in that environment, allowing us to gradually avoid poor decisions. When co-culturing heteroplasmy cells with several chemical inducers, the sub-lethal kanamycin selection will particularly affect cells with the highest red fluorescence. These cells’ ability to adapt by lowering their fluorescence can impact decisions made based on highest fluorescence levels. To showcase the power of this strategy, we can use it in board games where there are no more than nine options per round. We map chemical inducers to the opponent’s moves to determine the bacteria’s decision (Fig. 3b). Multiple AdaptoCells co-cultures can serve as the bacterial player, playing against a trainer. The trainer makes selected moves, creating chemical patterns that induce the strains in the AdaptoCells populations, leading to the bacteria responding with their F-weights, with the highest weight signaling the next bacterial move. After a loss, we apply kanamycin adaptation to every AdaptoCell involved in the decision, considering the environment produced by the trainer’s positions in each round. Over multiple lost matches, the AdaptoCells’ weights decrease, avoiding repeating losing moves. This results in the bacteria never losing a match, regardless of the opponent’s plays.

### AdaptoCells adapt to play the tic-tac-toe game

We have chosen the common tic-tac-toe game to demonstrate a complex board game. The goal of the game is for either player to be the first to line up three of their symbol "X" or "O" in a row, column, or diagonal on the 3×3 board. Previous studies approached this game with DNA computing and 23 custom 3-input logic gates^21, 44^, but these works did not involve living cells nor the capability to learn gameplaying. In our study, we assigned the first player as the trainer (player X) and player O as a bacterial player, consisting of a set of AdaptoCells for each of the board positions (excluding the center, which is played first by player X). We assigned a chemical inducer to identify each of the 9 board positions of player X (center Fig. 3c). The cells play a match by measuring their F-weights through red and green fluorescence measurements (Methods and Supplementary Text). To determine O’s move, we induce all co-cultures at unoccupied positions with the inducers assigned to X’s played positions (experimental workflow in Fig. 4c, Methods and Supplementary Text). The co-culture with the highest F-weight decides O’s next move. If O loses the match, we apply a kanamycin adaptation operation to the AdaptoCells only at the positions occupied by O (Fig. 3b). The next matches are played with the updated co-cultures.

To measure the overall skill level of the game, similar to the Elo ranking^45^, we run computer simulations of all possible games using the measured F-weights. These simulations, parametrized with our experimental data (Supplementary Text section 3.1, Table S2), evaluate the percentage of wins or draws (referred to as expertise) in all possible matches.

An illustration of a tic-tac-toe match is shown in Figure 3c. Player X starts at the center (Round 0) and player O can move to any of the 8 remaining positions. The F-weights of the co-cultures at those positions are measured by inducing them with the inducers assigned to X’s move (3-oxohexanoyl-homoserine lactone, OC6, inducer assigned to 5, the center position). The position with the highest F-weight is selected as O’s move. In the next round, X makes another move and O’s move is determined by measuring the F-weights of the co-cultures at unoccupied positions after inducing them with both X’s moves. O loses in Round 2, and a kanamycin adaptation operation is applied to the AdaptoCell populations at positions occupied by O. This adaptation updates AdaptoCell populations at positions 4 and 8. The bacteria continue playing matches until a match ends in a draw and player O demonstrates mastery.

### AdaptoCells playing and adaptation Tic-Tac-Toe against one another

Players have AdaptoCell libraries of specific weights in cell cultures assigned to board positions (Fig. 4a), implementing the decision tree in Fig. 4e. We initially choose all cultures to have similar AdaptoCell weights (Fig. 4a). We performed a series of matches between the bacterial cultures (experimental workflow in Fig. 3c, Methods and Supplementary Text). To simplify the decision tree, at each round we allowed a maximum of 3 available positions (marked with blue dots) for each player (Fig. 4d).

After an opponent’s move, we induce all permissible positions with the associated chemical input. The position with the highest F-weight is deemed the AdaptoCell’s choice (Fig. 3a bottom). If maximal fluorescence values are similar (within one standard deviation), we randomly select the position (Fig. 4d, blue bars), ensuring decision tree exploration. The same results come from a single biological replicate, where the highest value dictates the decision, incorporating inherent biological randomness.

Expertise was assessed after each adaptation iteration by running simulations for all possible matches using the weights of the library measured from plate-reader fluorescence (Fig. 3f). To assess the adaptation capability of our AdaptoCell libraries, we simulated the libraries under a cycle of 10,000 randomised adaptations (Fig. 1g, Supplementary Text). Our experimental kanamycin adaptation allowed Player 2 to achieve mastery in only one adaptation more than the minimal rate of 2 adaptations (Extended Data Fig. 10c).

### AdaptoCells can master arbitrary 3×3 board games

Next, we asked if our AdaptoCell libraries would be general enough to accommodate other well-known games such as chess, where pieces can be captured. A very simple variant is Hexapawn^46^, which is played on a 3 x 3 board, with three chess pawns on each side. To further simplify the decision tree, we added the rule where pawns must capture if possible (Fig. 5b). We therefore designed custom AdaptoCell libraries for each bacterial player (Fig. 5a), implementing the decision tree in Fig. 5d. We ran a tournament until Player 1 reached mastery (Fig. 5c, Supplementary Text). The expertise increased after each adaptation iteration (Fig. 5f). Our experimental kanamycin adaptation only requires one extra adaptation over the best strategy (which we validated in a simulation of 10,000 randomised adaptations, Supplementary Text).

We now asked how much we could push our libraries to learn more complex game strategies experimentally. We considered a simplified variant of the game Go (also called Atari Go) in a 3×3 board, where the first capture ends the game^47^. We imposed a toroidal symmetry, where positions at the opposite sides of the board are considered adjacent (Extended Data Fig. 7, 8, 9) and we only allowed at each round a maximum of 3 available positions. A tournament with kanamycin adaptation for both bacterial libraries (Extended Data Fig. 7a) allowed reaching mastery in 5 adaptations (Extended Data Fig. 7b-c, Extended Data Figs. 8, 9). To further explore the limits of our 9YES library to learn new games, we used a naive 9YES library of 0.9 weights to achieve mastery, after successively playing adaptation tournaments of 3 and 20 cycles as first and second player of the full tic-tac-toe game, respectively (Extended Fig. 10d). Afterwards, the resulting library also achieved mastery at the toroidal Atari Go game after playing 1 and 6 adaptation cycles in the role of first and then second player (Extended Data Fig. 10e). Interestingly, the library was able to simultaneously master both games as first and second player (Extended Data Fig. 10f). When the resulting library was challenged to learn the most difficult game we sampled (we call 33L), it lost mastery at the previous two games (the second player expertise dropped to random levels, Extended Data Fig. 10g).

Our libraries reached mastery against non-expert players; playing against experts could accelerate adaptation through increased losses. We anticipate the weights could diminish for highly challenging games, as experienced when mastering 33L after the simultaneous mastery of tic-tac-toe and toroidal Atari Go. A solution could involve an operation on all AdaptoCells, called AdaptoCell fusion, which could conserve the expertise while boosting the small (non-vanishing) weights (Extended Data Fig. 10, Supplementary Text).

## Discussion

In summary, we have developed a strategy to enable gene circuits to rapidly adapt to new behaviours by replacing designated genetic elements with AdaptoCells. AdaptoCells produce a gene expression that linearly correlates with their weights (Extended Data Fig. 4), which allows for the construction of gene circuits with predefined behaviour. For instance, using a AdaptoCell of weight 0.5 will be equivalent to using a traditional gene circuit with an engineered promoter with half of its transcription rate. Currently, the engineering of promoters of a targeted transcription rate requires screening mutant libraries^48, 49^. The AdaptoCells could be generalised in multicellular computing to arbitrary gene circuits by permitting the replacement of all genes in a decision circuit with AdaptoCells.

Our AdaptoCell system offers a unique solution to the limitations faced in traditional quantitative control over gene expression levels. Rather than offering direct control, our system allows for simultaneous relative adjustments to all the induced sensor strains’ weights through the presence of chemical inducers and sub-lethal kanamycin. The weights of each sensor strain are linked to fluorescence, providing immediate feedback to the experimenter on the changes made. This enables a progressive adaptation towards a desired behavior without the need for prior knowledge of the optimal weights or manual weight setting.

Examining the capabilities of AdaptoCells as a single-layer artificial neural network (ANN)^50, 51^ or perceptron reveals valuable insights. The tic-tac-toe AdaptoCell libraries associated to each board position would correspond to a network of eight 9-input neurons and 72 synaptic weights. As ANNs with positive weights and decisions based on the highest output signal can approximate any function^52^, we anticipate that the AdaptoCells could learn many games on a 3×3 board. Unlike traditional backpropagation algorithms^53^, our adaptation method does not require knowledge of the weights or computation of the prediction error, making AdaptoCell adaptation more closely mimic the functioning of the brain. This "training" is achieved through repeated rounds of kanamycin adaptation, using chemical patterns and red fluorescence protein expression levels as input and prediction output. Incorrect decisions trigger a decrease in the weights of the responsible AdaptoCells, leading to gradual improvement and optimized decision-making. In the same way that a perceptron can be extended to multilayer ANNs, AdaptoCells can be extended to higher orders through cell-cell communication, like other multicellular circuits^3, 4^ (box I).

The improved adaptation in our extensions of the 9YES library to more complex promoter regulations, combinatorial (4AND library) or hybrid (3HYB), suggests that by extending our gene circuits (Box 1) to include more complex regulations would enhance the adaptation capability. Moreover, the use of libraries of hybrid AdaptoCells together with the use of appropriate selections (Box 1) provides a general way to adapt the topology of gene circuits^54^ (Extended Data Fig. 2). The 9YES library could also learn all the 1,500 games we tested, and it could even simultaneously master more than one game (Extended Data Fig. 10), showing that the information stored in the weights was still enough to learn new different algorithms. This highlights the aptitude of 9YES for general-purpose adaptation. But, when pushing the 9YES library towards its limits, we found that it was not able to learn the 33L game without losing the mastery at toroidal Atari Go, suggesting that we possibly reached its adaptation capacity.

As the adaptation of gameplaying includes as a particular case the adaptation of the input patterns associated to a winning game, AdaptoCells may be used in applications such as pattern discrimination problems in therapeutics^55^, where a reengineered microbiome is trained to adapt to certain chemical patterns, triggering a specific therapeutic delivery. AdaptoCells preserved their weight through growth cycles alternating with stationary states, both in liquid and solid cultures over several days (Extended Fig. 1, 3b,d), implying potential use in ecosystem-level gene circuits^56^. AdaptoCells could also be used with alternative selection markers, in single-cell circuits, or to explore alternative circuit topologies (Box 1). We could foresee adding an extra AdaptoCell library to each player that would receive the output of the first library through a cell-cell communication system (Box 1), mimicking a hidden layer in a neural network. This would enable the processing of more complex information and, therefore, adaptation more advanced games. Further developments could involve a way to genetically encode the computation of the maximum output among positions^17^, as well as research into the possibility of naturally occurring AdaptoCells in prokaryotic or eukaryotic systems as a non-Darwinian adaptation tool^57^ (Box 1). Kanamycin adaptation with AdaptoCells provides a strategy for the unsupervised adaptation of complex gene circuits with a large, unknown number of interactions, which will allow for the engineering of genetically-encoded general-purpose computational devices capable of self-adaptation.

## Material and methods

### Strains and growth media

Cells were routinely grown aerobically at 37 °C in 14-ml tubes (Supplier Scientific Laboratory Supplies Ltd, UK) in LB (1% tryptone, 0.5% yeast extract and 1% NaCl), LB agar plates (1.5% Select agar, Sigma Aldrich), or M9 medium (1x M9 salt (Sigma Aldrich), 100 μM CaCl2, 2 mM MgSO_4_, 10 μM FeSO_4_, glycerol 0.8% v/v) supplemented with casamino acids 0.2% w/v, 1 μg/mL thiamine, 20 μg/mL uracil, 30 μg/mL leucine, using NaOH to adjust the pH to 7.4. The base strain for all cloning experiments was *Escherichia coli* strain Top10 (genotype [F-mcrAΔ(mrr-hsdRMS-mcrBC) Φ80lacZΔM15ΔlacX74 recA1 endA1 araΔ139 Δ (ara, leu)7697 galU galK λ-rpsL (StrR) nupG]), which was grown in LB medium at 37 °C and 200 rpm with antibiotics. The default antibiotic concentrations used are: carbenicillin, 80 μg/mL; kanamycin 50 μg/mL; ampicillin, 100 μg/mL; chloramphenicol, 34 μg/mL. When recovering a culture from frozen glycerol stock, bacteria were inoculated into 2ml LB medium and grown overnight at 37 °C, 200 rpm with ampicillin (100 µg/ml). The chloramphenicol resistance gene from the Marionette DH10B strain was cured (eliminated) by electroporating 60 ng pE-FLP plasmid^58^ into their competent cells and selecting on 50 μg/mL ampicillin agar plate at 30 °C for one day. This cured strain was used for all AdaptoCell adaptations and characterisations.

### Plasmid construction

All the plasmids were generated Gibson assembly^59^ by relying on gene synthesis and PCR using Q5 master mix DNA Polymerase (New England Biolabs, USA) with 40-bp homology domains. DNA was purchased from Integrated DNA Technologies (USA). The plasmids P1 and P2 were designed to have the same size (4,774 bp in total) and a common backbone consisting of: origin of replication (medium copy number pMB1), ampicillin resistance cassette (from the pSEVA191 plasmid^60^) and the spy terminator^61^ to isolate the promoter. The replicon pMB1 (derivative of ColE1) was amplified from the plasmid pWT018e (Addgene #107886)^28^ by introducing a mutation in the RNAII gene (T to C) to lower the copy number^62, 63^. To create the AdaptoCell libraries, a promoter-specific spacer followed by the promoter were inserted into the common backbone plasmid of both P1 and P2 plasmids (except for the hybrid AdaptoCell of Fig. 1i, where each plasmid has a different promoter). A terminator was introduced when the plasmid size restriction allowed for it (Supplementary Information). The promoter was followed by a promoter-specific spacer and by a RiboJ ribozyme insulator^64^ used to insulate the downstream transcript. The RiboJ was followed by a weak ribosome-binding site (BBa_B0064 from the Registry of Parts) and by the mCherry (from plasmid from PAJ310^65^, with a C-terminus addition of a Gly-Ser-Gly amino acid sequence to match the size of EGFP) and EGFP (from plasmid pWT018a) genes for the P1 and P2 plasmids respectively. After the fluorescent protein genes, the antibiotic resistance cassette CmR-dKanR and dCmR-KanR from pWT018e and pWT018a was added polycistronically (using the ribosome-binding site BBa_B0064) to the plasmids P1 and P2 respectively. CmR-dKanR contains a chloramphenicol resistance gene fused (using a Gly-Ser-Gly-Ser-Gly-Ser linker) to a kanamycin resistance gene with a Asp208Ala amino acid mutation to obliterate its activity. dCmR-KanR contains a chloramphenicol resistance gene with a His195Ala amino acid mutation to obliterate its activity (and an Ala29Thr amino acid mutation that appeared during our cloning) fused to a kanamycin resistance gene using a Gly-Ser-Gly-Ser-Gly-Ser linker.

### Combinatorial promoter engineering

Our approach to engineering combinatorial promoters involved integrating transcription factor operator sites into existing promoters, aiming to trigger expression only when two specific chemical inducers reached sufficient concentrations (AND logic gate on the inducers). This precisely tuned response was achieved by incorporating operator sites from different transcription factors into various promoters from the Marionette library at exact locations, while ensuring no interference with -10 and -35 sequences and those upstream of -35. Beginning with the pTac-Tet promoter, we incorporated a 19 base pair (bp) tetO2 operator. The pTac-Van promoter was enhanced by the addition of a 30 bp VanO2 operator, as described by Gitzinger et al. in 2012, differing from the operator previously used in the Marionette study. Similarly, this modified VanO2 operator was also introduced into the pTet-Van promoter. For the pTet-Bet promoter, we designed a novel 22 bp BetO operator, inspired by the work of Stanton et al. 2013. The pTet-Ttg promoter was modified by integrating a 30 bp TtgR operator from Krell et al. 2007, which deviates from the operator in the Marionette paper by two bases. We inserted a 30 bp PhlF operator from Abbas et al. 2002 into the pTet-PhlF promoter, placed in reverse orientation to avoid potential interference with a -10 sequence. The pVan-Ttg and pVan-Tac promoters were equipped with the TtgR operator from Krell et al. 2007 and the symmetric version of the lacOsym operator from Lutz et al. 1997, respectively. Similarly, the symmetric lacOsym operator was included in the pBet-Tac, pTtg-Tac, and pPhlF-Tac promoters. For the 4ANDMEM library, we engineered the combinatorial promoters pTet-Ttg, pTac-Van, pVan-Ttg, and pTac-Tet by embedding operators for TtgR^66^, VanR^67^, TtgR^66^ and TetR^68^ downstream of the pTet, pTac, pVanCC, and pTac promoters^68^ at positions +1, +20, +5, and +20, respectively. These operators spanned lengths of 30 bp, 30 bp, 30 bp, and 19 bp. The TtgR operator was oriented in reverse to prevent the formation of a tandem promoter, owing to its inherent -35 and partial -10 sequences.

### AdaptoCell transformation

To create a cell carrying a AdaptoCell, the P1 and P2 plasmids were co-transformed at equimolar concentrations (80 ng each) in the cured *E. coli* Marionette DH10B strain. AdaptoCell selection was achieved on LB agar plates with ampicillin (60 μg/ML), kanamycin (17.5 μg/mL), chloramphenicol (11 μg/mL) and cognate chemical inducers for the promoter(s) in the AdaptoCells. The concentrations for the chemical inducers (pSal: Sal, Sodium salicylate; pTet: aTc, Anhydrotetracycline HCl; pBetI:Cho, Choline chloride; pBAD: Ara, L-Arabinose; pLux: OC6, 3OC6-AHL; pCin: OC14, 3OHC14:1-AHL; pTtg: Nar, Naringenin; pVan: Van, Vanillic acid; pTac: IPTG, Isopropyl-beta-D-thiogalactoside) are: pSal (Sal 60 μM), pTet (aTc 0.1 μM), pBetI (Cho 6,000 μM), pBAD (Ara 2,000 μM), pLux (OC6 6 μM), pCin (OHC14 5 μM), pTtg (Nar 250 μM), pVan (Van 40 μM), pTac (IPTG 250 μM), pTet-Ttg (aTc 0.1 μM, Nar 250 μM), pVan-Ttg (Van 25 μM, Nar 250 μM), pTac-pTet (IPTG 250 μM, aTc 0.1 μM), pTac-Van (IPTG 250 μM, Van 25 μM). When P1 and P2 had different promoters (Fig. 1i), they were both induced with their cognate chemicals. Single colonies were picked and grown overnight with ampicillin (100 µg/ml). On the next day glycerol stocks were made and stored them at -80 °C until use.

### AdaptoCell stability

Glycerol stocks of bacteria carrying AdaptoCells were inoculated and grown them overnight with ampicillin (100 µg/ml) in either liquid or solid LB medium. The following day, a sample was taken from the culture to make glycerol stock and stored at -80 °C while the main culture continued to grow. We repeated this process for 5 days.

### AdaptoCell adaptation

Glycerol stocks of bacteria carrying AdaptoCells were inoculated and grown them overnight with ampicillin (100 µg/ml) after which the cultures were diluted 1:200 into freshly prepared 1.5mL M9 medium with carbenicillin, cognate inducer and custom kanamycin concentrations for each AdaptoCell: pSal 1.5 μg/mL, pTet 2.5 μg/mL, pBet 4 μg/mL, pBAD 2.5 μg/mL, pLux 1 μg/mL, pCin 1.5 μg/mL, pTtg 1.5 μg/mL, pVan 3.5 μg/mL, pTac 4 μg/mL, <u>pTet</u>-<u>Ttg</u> 1.5 μg/mL, <u>pVan</u>-<u>Ttg</u> 1.5 μg/mL, <u>pTac</u>-<u>Tet</u> 1.5 μg/mL, <u>pTac</u>-<u>Van</u> 1.5 μg/mL. The chemical inducers concentrations used for each AdaptoCell are: pSal (Sal 100 μM), pTet (aTc 0.2 μM), pBetI (Cho 10,000 μM), pBAD (Ara 4,000 μM), pLux (OC6 10 μM), pCin (OH14 7 μM), pTtg (Nar 1,000 μM), pVan (Van 100 μM), pTac (IPTG 500 μM), pTet-Ttg (aTc 0.2 μM, Nar 1,000 μM), pVan-Ttg (Van 100 μM, Nar 1,000 μM), pTac-pTet (IPTG 500 μM, aTc 200 μM), pTac-pVan (IPTG 500 μM, Van 100 μM). After refreshment the culture was split into three individual cultures which were then carried separately in subsequent experiments. The resulting culture tubes were incubated for 8 hours at 37 °C with shaking (200 rpm). Finally, glycerol stocks were made and stored at -80 °C. This process was repeated 4 or 5 times for each AdaptoCell. For the AdaptoCell adaptation in distributed multi-cellular co-cultures, glycerol stocks of bacteria with AdaptoCells were inoculated and grown separately overnight with ampicillin (100 µg/ml). The following day, cultures of bacteria with different mermegulons were mixed in equal volumes, then the resulting mixture was split in two parts, one made into glycerol stock and stored at -80 °C and the other part was used for adaptation. The culture for adaptation was diluted 1:200 in LB and grown for 8 hours in presence of carbenicillin, with the cognate inducer(s) and the specific kanamycin concentration.

### AdaptoCell fusion

This operation merges two cultures of the same AdaptoCell but different weight. If a AdaptoCell of weight 0.5 is added to all the members of a library, the weights will increase if lower than 0.5. However, the expertise of the library (for any game) remains unchanged. Two glycerol stocks of bacteria carrying the same AdaptoCell of different weights were inoculated and grown overnight with ampicillin (100 µg/ml). The cultures were grown independently and before characterisation 200 µL of each were mixed and 200 µL was used to make glycerol stock and stored it at -80 °C.

### AdaptoCell characterisation using fluorescence plate reader

Glycerol stocks of bacteria carrying AdaptoCells were inoculated and grown overnight with ampicillin (100 µg/ml). The overnight culture was refreshed by diluting 1:2000 in freshly prepared M9 medium with cognate inducer(s) and carbenicillin (80 μg/mL). The concentrations for the promoter’s chemical inducers were: pSal (Sal 100 μM), pTet (aTc 0.2 μM), pBetI (Cho 10 mM), pBAD (Ara 4 mM), pLux (OC6 10 μM), pCin (OH14 7μM), pTtg (Nar 1 mM), pVan (Van 100 μM), pTac (IPTG 500 μM), pTet-Ttg (aTc 0.2 μM, Nar 280 μM), pVan-Ttg (Van 25 μm, Nar 280 μM), pTac-Tet (IPTG 500 μM, aTc 200 μM), pTac-Van (IPTG 500 μM, Van 21 μM). Following an additional 3 hours of growth, 200 μL samples of each culture were added to a 96-well plate (Custom Corning Costar) with both technical and biological replicates. The plate was loaded in an Infinite F500 microplate reader (Tecan) at 37 °C with shaking. OD600 was assayed with an automatic repeating protocol of absorbance measurements (600 nm absorbance filter) and fluorescence measurements (465/35 nm excitation filter––530/25 nm emission filter for EGFP and 580/20nm excitation filter -635/35 nm filter for mCherry) were taken every 15 min during 18 hours of growth. The F-weight (W), which agrees with the weight for a monoculture, was 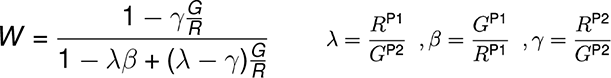 estimated by using:

where and R and G are the red and green fluorescence values of an AdaptoCell culture or co-culture. R^P1^ and R^P2^ are the red fluorescence value of fully induced P1- and P2-only cells and G^12^ and G^P2^ are the green fluorescence value of fully induced P1- and P2-only cells. We approximated γ = 0. The λ and β values were computed for each promoter and are included in Table S19. For the fluorescence characterisation of distributed multi-cellular co-cultures, only one AdaptoCell was induced at a time and the weight of the mixed culture was multiplied by the number of component strains.

### Fluorescence plate reader data analysis

Excel (Microsoft) with bespoke macros was used for data analysis (Supplementary Information). In summary, OD600 values were synchronised at OD600 = 0.2, where linear interpolation was used to infer fluorescence (F) and OD600 values at non-measured times. The growth rate was fitted against an exponential and poor fits (r^2^<0.9) were discarded. The fluorescence reported was computed as a constant slope for a non-linear fitting of the F to the OD600. The weight was calculated by the non-linear fitting of the whole dynamics of the red fluorescence of the strain of interest against the red fluorescence of P1-only cells averaged among biological replicates.

### AdaptoCell characterisation using flow cytometry

Glycerol stocks of bacteria carrying AdaptoCells were inoculated and grown overnight with ampicillin (100 µg/mL). On the next day the cultures were refreshed with 200 μl dilution in 2 mL LB with carbenicillin (80 μg/mL) and cognate inducer, and later incubated for 4 hours at 37 °C with 200 rpm shaking. The concentrations for the promoter’s chemical inducers were the same used for the plate reader assay. After 4 hours of growth, when the log phase was reached, the cultures were centrifuged at 3000 rpm for 10 minutes at 4 °C. The pellets were resuspended in 2 mL PBS solution and 200 μL samples were loaded in 96 well plate. The plates containing cell cultures were processed using a BD LSRFortessa High Throughput Sampler (HTS) with the following settings: at a Sample Flow Rate of 1ul/second; Sample Volume 5 μl; Mixing Volume 50 μl; Mixing Speed 200 μl/sec; Number of Mixes 5; Wash Volume 200 μl; BLR Enabled. A maximum of 50,000 events were recorded for each sample.

### Flow cytometry data analysis

The weights were calculated with the equation (see Supplementary Information for its derivation) weight = 1/(1 + 𝜆(G/R − 𝛽)), where 𝜆 = 𝑅*_p_*_1_/𝐺*_p_*_2_, 𝛽 = 𝐺*_p_*_1_/𝑅*_p_*_1_. *R_P1_* and *G_P1_* are the mean red and green fluorescence of P1-only cells. We computed lambda and beta for each promoter separately. Weight values higher than 1 or lower than 0 were removed. We calculated the weight using the red and green fluorescence values of the strain of interest. To predict a AdaptoCell’s weight after a given number of adaptation iterations, the 2% of cells with the highest red fluorescence were recursively pruned followed by re-normalising the area of the distribution (Extended Data Fig. 5b, Supplementary Text).

### AdaptoCell characterisation using Sanger sequencing

Glycerol stocks of bacteria carrying AdaptoCells were inoculated and grown overnight with ampicillin (100 µg/ml). The plasmid DNA (GeneJET, Thermofisher) was purified for Sanger sequencing. Minipreps from single-strain bacterial colonies carrying AdaptoCells were sequenced with the CamR reverse primer annealing to a common sequence of P1 and P2. For distributed multicellular co-cultures, forward primers specific to only one of the promoter regions of the AdaptoCells found in the miniprep were used. All primers are included in Supplementary Table 5. Minipreps of the P1- and P2-only cells were also sequenced with the CamR primer. The mixed read chromatogram traces (AB1 files) from the Sanger sequencing (GATC, Germany) were kept for subsequent analysis.

### Sanger sequencing data analysis

To compute the weight, the mixed reads’ chromatogram traces from the Sanger’s AB1 files were analysed by aligning them against the traces of P1- and P2-only cells using a custom algorithm (Supplementary Information) implemented in Biopython^69^ (a recently published method^70^ did not give adequate results for AdaptoCells). In summary, our algorithm firstly aligns the chromatogram traces by basecalling them and performing a sequence alignment against EGFP-mCherry. Afterwards, it estimates the weight from the average ratio of the mixed read’s peaks against the P1- and P2-only cells’ peaks at EGFP-mCherry mismatch positions. The EGFP-mCherry match positions were used to normalise the traces.

### AdaptoCell weight characterisation by qPCR

Glycerol stocks of bacteria carrying AdaptoCells were inoculated and grown overnight with ampicillin (100 µg/ml). Plasmid DNA was extracted using GeneJET Plasmid Miniprep Kit (Thermo ScientificTM, UK K0503) and quantified using the NanoDrop® ND-1000 UV-Vis Spectrophotometer. Quantitative real-time– PCRs (qRT–PCR) analyses were performed on the StepOnePlus™ Real-Time PCR System (Thermo ScientificTM) using the Fast SYBR-Green master mix (Thermo ScientificTM, UK K0503) and following the manufacturer’s instructions. The P1 plasmid was quantified using the RG1083 and RG1084 primers specific for the mCherry coding sequence. The P2 plasmid was quantified using the RG1087 and RG1088 primers specific for the EGFP coding sequence. The RG1080 and RG1082 primers which anneal to a common sequence of P1 and P2 construct backbone were used as endogenous control for the relative quantification of P1/P2 plasmids. All primers are included in the Supplementary methods.

### AdaptoCell fluorescence characterization in solid medium

We surrounded the cocultures on the agar plate with aluminum foil masks and transferred them to a new LB agar plate using a standard replica plating procedure. Both the original and replica plates were incubated at 37 °C for 1 hour, and the original plate was further incubated for 6 hours to restore the cocultures. We added 0.5 µl of the necessary inducers to the cocultures on the replica plate, and let the cells grow for 6 hours at 37 °C. After that, we stored them overnight at 4 °C. To image the cocultures for fluorescence expression, we used a blue LED (470 nm) transilluminator with an amber filter unit (580 nm).

### Experimental strategy for gameplaying with AdaptoCell libraries in liquid medium

Bacterial cultures carrying AdaptoCells are assigned to each board position for each bacterial player according to a specific game and decision tree. For positions with more than one AdaptoCell, the cultures are grown individually or together as an equal volume co-culture. To encourage both bacterial players to take random decisions before any adaptation, AdaptoCells of equal weights within one standard deviation in their biological replicates are initially chosen. Player 1 makes the first move. For tic-tac-toe and toroidal Atari Go, Player 1 initially plays in position 5. Moves are determined by the highest normalised red fluorescence in plate reader across unoccupied (and allowed, in case of games with simplified decision trees) positions after growing from glycerol the AdaptoCell cultures and inducing them with the chemicals assigned to the opponent’s positions. When positions contain more than one culture, all the normalised red fluorescence values of those cultures are added. If the highest red fluorescence involves more than one position (within one standard deviation in the biological replicates) then we choose randomly among these positions. This cycle is repeated until one of the ending rules is matched. At the end of each game, we assessed if any of the two players has lost, case in which kanamycin adaptation was applied to the loser. For this, AdaptoCell adaptation was conducted for each move played by adding the chemicals of the opponent’s positions and kanamycin. The glycerol stocks were updated and the matches continued until both players reached mastery.

### Experimental strategy for gameplaying with AdaptoCell libraries in solid medium

The game is initiated by growing the AdaptoCell cultures, then distributing them across all positions of a game board-like agar plate except the center. This plate, referred to as the parental plate, which is only measured by replica-plating. The game proceeds in rounds where inducer assigned to the positions occupied by player X, is added to all positions. After replica-plating the parental plate, a period of incubation and cooling, the cultures are imaged for fluorescence expression. The brightness of fluorescence is quantified using the equation for the F-weight (eq. 2.1 in Supplementary Text), and the F-weight culture is considered player O’s move. This process is repeated for each round until a winning condition, or end rule, is met (see details of the protocol in the Supplementary Text). If the bacteria lose the game, they undergo an adaptation process, where we do a replica plating of the parental plate, where the cocultures are exposed to inducers that represented the player X’s moves in the rounds of the match, and kanamycin. This is performed in a given incubation period (see Supplementary Text). Then, another plating procedure is performed to halt the adaptation process and to remove the inducers and kanamycin. After an overnight storage, a new parental plate with the newly adapted bacteria is ready for the next game. The glycerol stocks are made and updated and the matches continued until both players reached mastery.

### Computer simulation of bacterial gameplaying

A C++ code was written to simulate the capability of bacteria carrying AdaptoCells to play and learn different games. The red fluorescence of a AdaptoCell is modelled by multiplying its weight to the promoter’s effective transcription rate. The effective transcription rate is fitted from our experimental data (supplementary Text). Mimicking the experimental gameplaying strategy, the move of a player is computed from the available position with highest fluorescence after having induced the AdaptoCell’s promoters with the chemicals corresponding to the opponent positions. Before the start of a tournament, both players are set as learners. If a player loses a game, kanamycin adaptation is applied by decreasing the weights of the AdaptoCells at the positions played by a pre-defined coefficient of 0.05. To evaluate the expertise of a given set of weights (corresponding to a AdaptoCell library), a player that plays randomly (uniform probability) among available positions was used. This player was used as a non-learner to calculate the expertise of the learners after each kanamycin adaptation by playing 10,000 games and checking the percentage of positive results. This computation of expertise was also used with an experimental AdaptoCell library once we have measured the weights by fluorescence or DNA quantifications. The successive adaptation tournaments of tic-tac-toe, toroidal Atari Go and 33L (Extended Fig. 10d) where done by playing each resulting 9YES library against a new 9YES library (with starting weights equal to 0.9) able to learn by kanamycin adaptation. The tic-tac-toe tournament after achieving mastery at 33L included a AdaptoCell fusion operation at match 9.

### Randomised game rules and adaptations

For Fig. 1d, we considered a 1,500-sample of all possible games where the pieces cannot be moved nor captured. We generated 10,000 games with 110 random zero-sum end-of-game rules, each specifying 3 or 4 occupied board positions defining winning or losing the match. To generate the adaptation tournaments used in Fig. 3g, Fig. 4g and Extended Fig. 8h, we uniformly selected at random either 3 (Hexapawn) or 4 (tic-tac-toe and toroidal Atari Go) AdaptoCells from the library of a player to undergo kanamycin adaptation. Examples are included in Extended Data Fig. 10c.

### Statistical analyses

All measurements were taken from distinct samples and the experiments were not randomised. All Pearson’s r coefficients were determined by linear regression analysis. Non-parametric ANOSIM tests done with the Vegan R package were used to assess the hypotheses of statistically significant differences between two bacterial cultures transformed with AdaptoCells. AdaptoCell stability (Fig.1b), relative activation (Fig. 1d,i), and the weight decrease produced by kanamycin adaptation (Fig. 3e) were tested.

### Reporting summary

Further information on research design is available in the Nature Research Reporting Summary linked to this paper.

### Data availability

All relevant data are included as Source Data and are available from the corresponding author on reasonable request. Strains and plasmids used in this study are available from the corresponding author on reasonable request and from the Addgene repository.

### Code availability

The software used for simulations and data analysis is available from the corresponding author on reasonable request.

## Acknowledgments

RG acknowledges the departmental allocation from the School of Life Sciences (Faculty of Natural Sciences, Keele University). MI acknowledges funding from BBSRC BB/P020615/1 (EVO-ENGINE) and the Volkswagen Foundation (LIFE: 93 065). AJ was funded by the Ministerio de Ciencia e Innovacion GeneCircuits++ PID2020-118436GB-I00 (GeneCircuits++), BBSRC BB/P020615/1 (EVO-ENGINE), EU grants EVOPROG 610730 and PhotoSynH2 HORIZON-EIC-2021-PATHFINDERCHALLENGES (grant agreement 101070948), EPSRC-BBSRC BB/M017982/1 (WISB centre), ONRG (BacMorse), and the departmental allocation from the School of Life Sciences (U. Warwick). We thank J. Penadés, M. Kushwaha and V. de Lorenzo for their comments.

---

**Extended Data Fig. 1.**
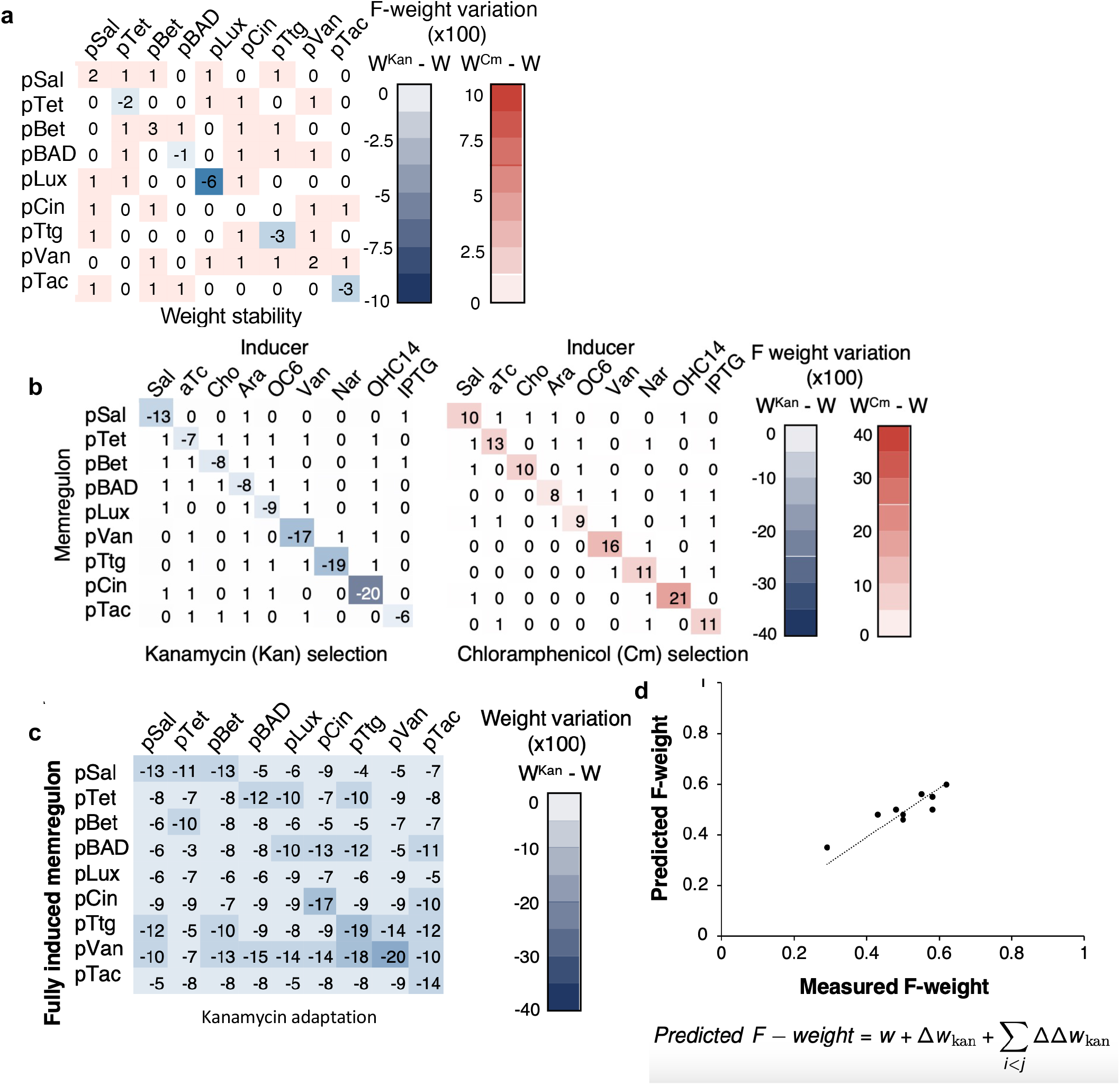
F-weights are stable in AdaptoCell mono-and co-cultures. **a,** After four days of daily replica plating, freezing and replica copying, the F-weight variation in co-cultures of two AdaptoCell strains demonstrates minimal and non-statistically significant change (lowest p>0.1). This data informs a pairwise model (refer to Supplementary Text and panel **d**). **b**, Adaptation of single-sensor AdaptoCell strains is activation-dependent. Statistically significant weight variation (p<0.01) occurs when strains grow on agar plates with the orthogonal inducer, under kanamycin or chloramphenicol selection. The inducer and antibiotic concentrations are detailed in the Methods. **c**, Composition of co-culture influences kanamycin adaptation. Co-cultures of two memregulon strains exhibit significant fluorescent weight variation (p<0.05) under kanamycin adaptation when only one component strain is fully induced. Culturing two AdaptoCell strains together results in less weight variation during kanamycin adaptation, as inactive promoter cells’ reduced growth artificially boosts the weight of active promoter cells. **d**, A pairwise model using the F-weight variation in co-cultures of two AdaptoCell strains (as done in panel **a**) can predict the F-weight variation in co-cultures of nine AdaptoCell strains (R2=0.91).

**Extended Data Fig. 2.**
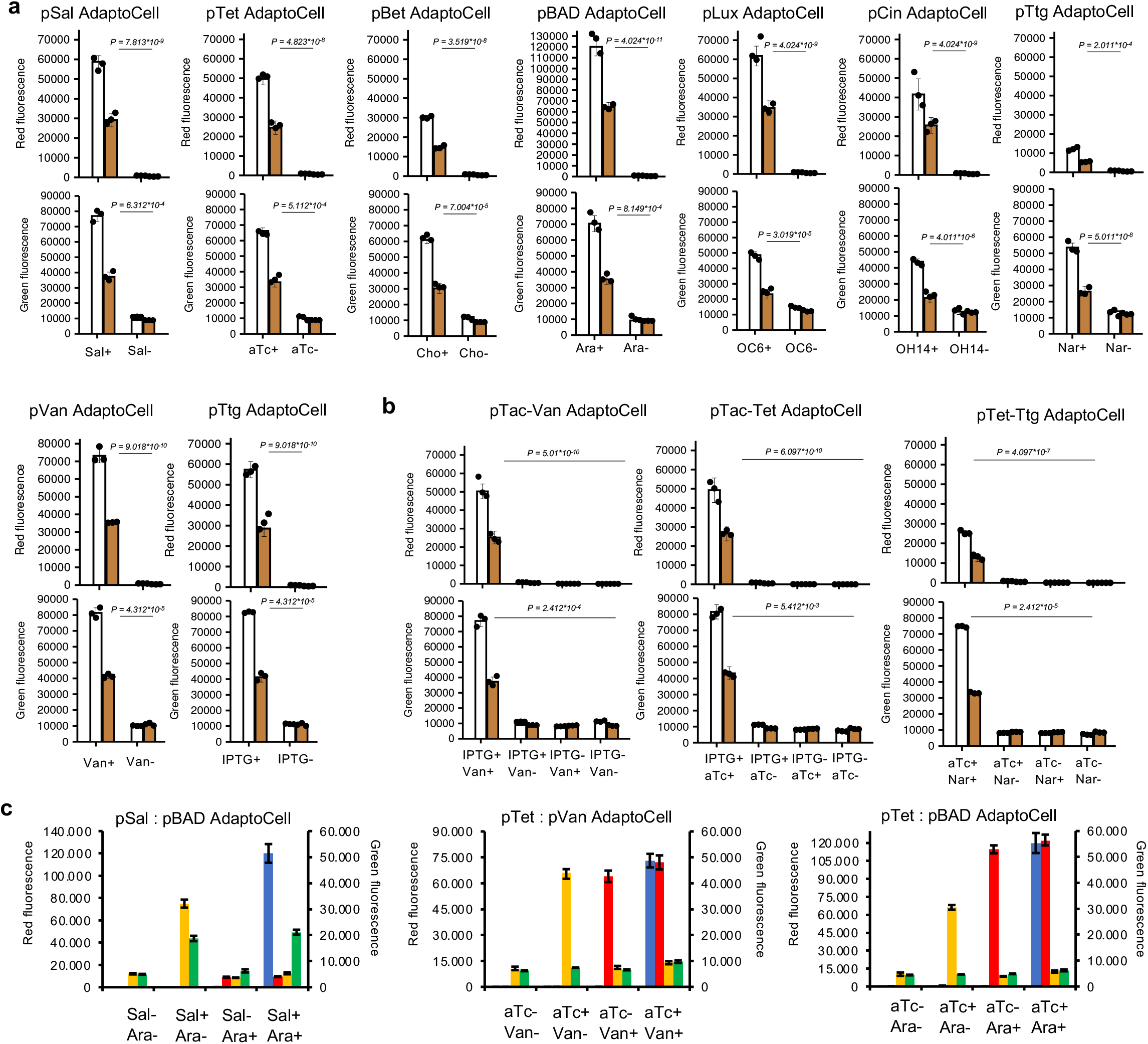
AdaptoCell computation capabilities. Red and green fluorescence of cells transformed with either P1 or P2 (white-filled bars) and 0.5 weight AdaptoCells (brown bars) grown in presence (at saturating levels, see Methods) or absence of cognate inducer for the 9YES AdaptoCell library (**a**) the 3 remaining 4AND library members (see Fig. 1), (**b**), and the 3HYB library (**c**). The 4AND and 3HYB libraries are grown in the presence or absence of each of the two cognate inducers.

**Extended Data Fig. 3.**
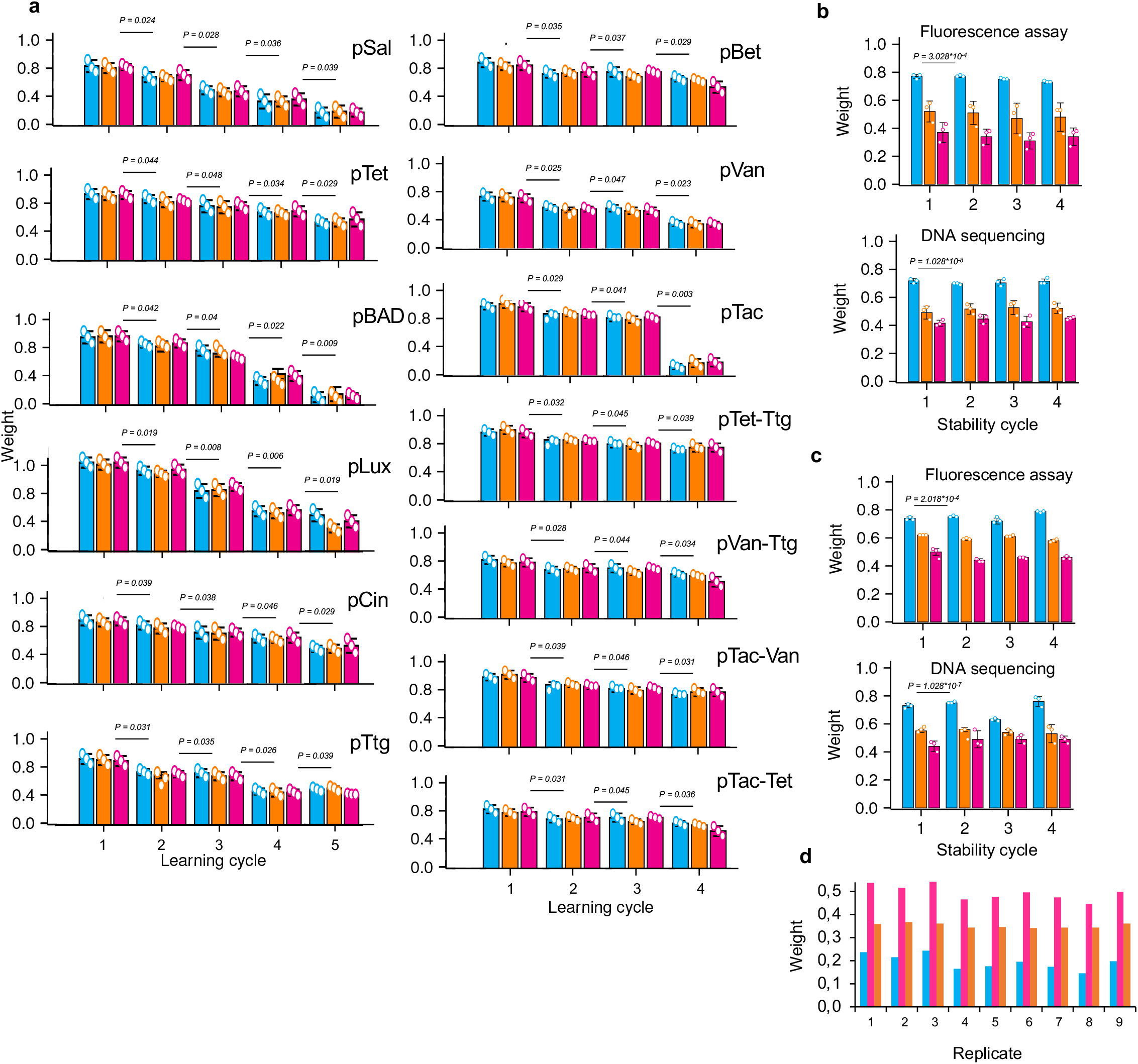
Adaptation and stability of AdaptoCell’s gene expression. **a,** Fluorescence analysis-based weights (F-weights) of 9YES and 4AND AdaptoCell libraries following five and four sequential adaptation experiments, respectively (for equivalent DNA sequencing plots, see Supplementary Information). Different colors denote different lineages. **b**, pBAD AdaptoCell weight, as measured through fluorescence plate reader assay and DNA sequencing, for three different AdaptoCells in stationary state liquid cultures at 37°C, without induction or adaptation. **c**, pBAD AdaptoCell weight in stationary state solid medium cultures at 37°C, absent induction or adaptation. **d**, Fusion operation characterization showing preservation of any player’s library expertise for a given game. We added an equal volume of the same AdaptoCell culture (pBAD with a 0.5 weight) to a given AdaptoCell culture (pBAD). Weight measurements, via ratiometric fluorescence assay, before (blue bars) and after a 1:1 co-culture (orange bars) with a pBAD AdaptoCell of weight 0.5 (red bars) are shown. AdaptoCell fusion (refer to Supplementary Text) helps maintain weights after numerous kanamycin adaptations, facilitating continued adaptation.

**Extended Data Fig. 4.**
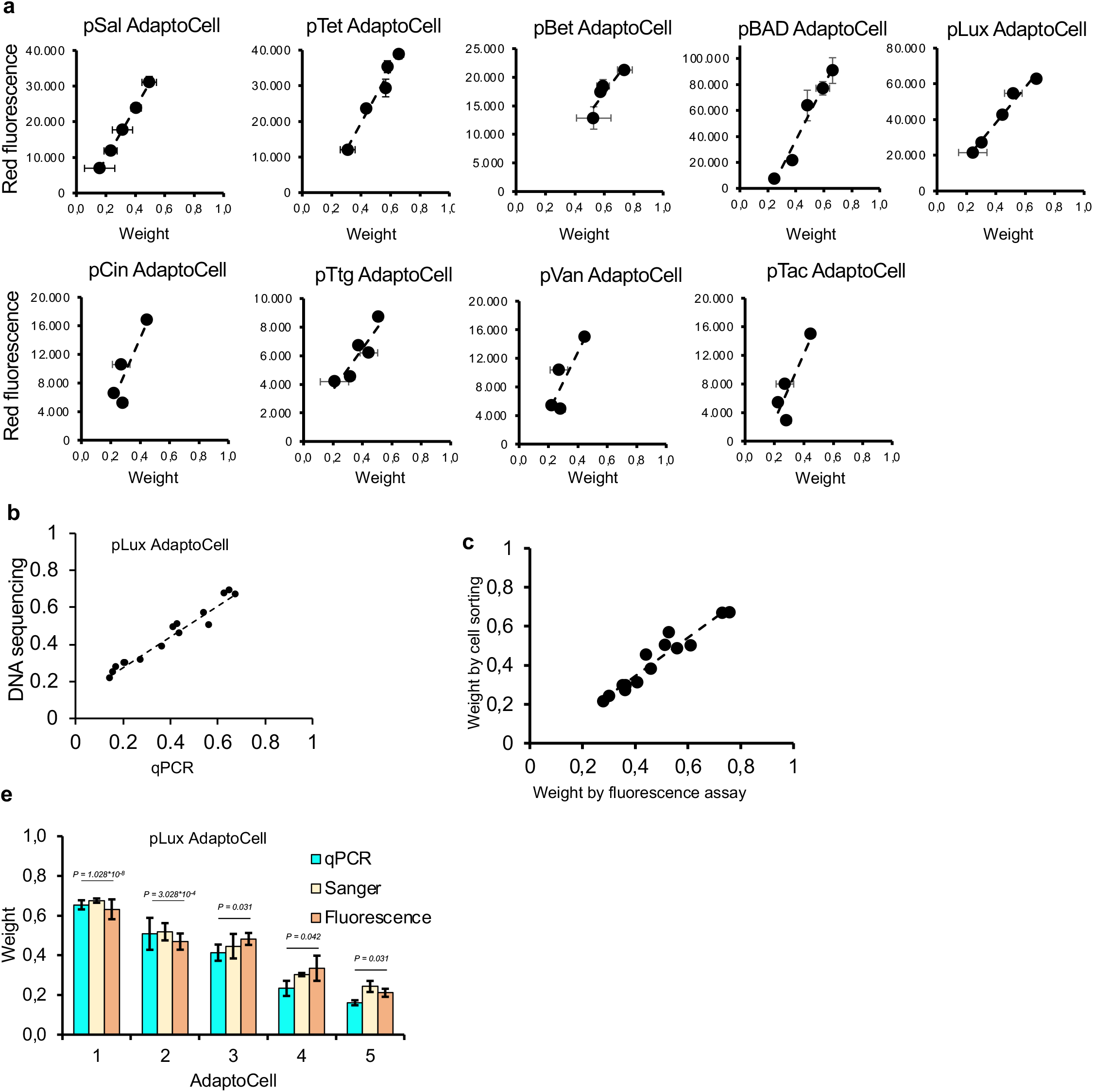
AdaptoCell’s weight measurement agree when using different techniques. **a**, AdaptoCells of different weights give a predictable gene expression (R^2^=0.99, (± s.d. for n = 3). **b,** Correlation between weights measured by Sanger sequencing against qPCR data. **c**, Correlation between weights measured by cell sorting against fluorescence assay for the pLux AdaptoCell. (R^2^=0.92) **e,** Example of simultaneous characterization of each round of adaptation of the pLux AdaptoCell using Sanger sequencing, qPCR and fluorescence assay data (averge ANOSIM R=0.82).

**Extended Data Fig. 5.**
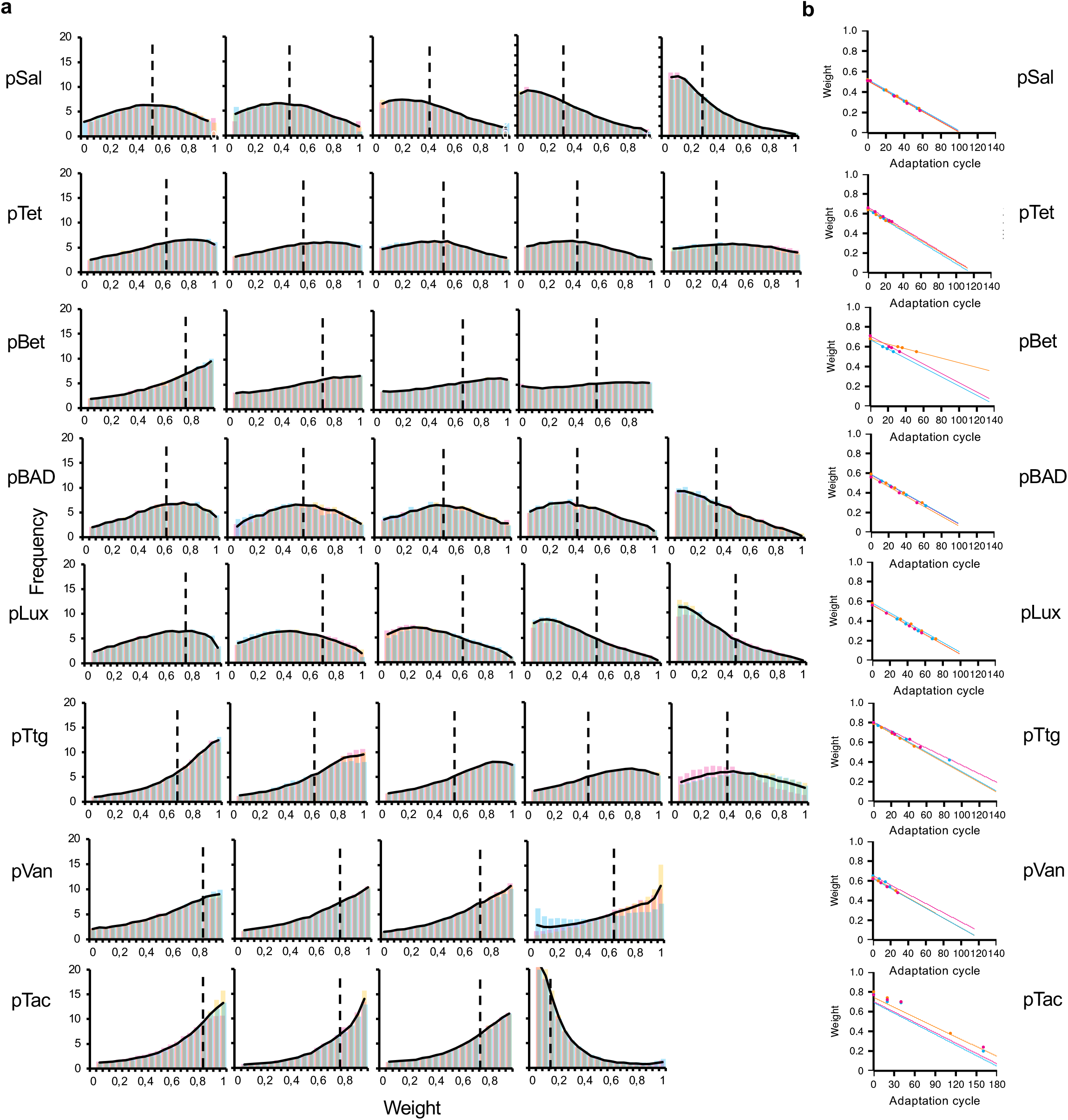
AdaptoCell Adaptation measured by flow cytometry. **a,** Single-cell weight distribution for eight representative AdaptoCells analyzed via flow cytometry (see Methods) after each adaptation cycle, presented in three biological replicates (blue, pink, and yellow bars). The solid black line illustrates the average model prediction value for three replicates, with the vertical dashed line indicating the mean value. **b**, Model-predicted weights originating from a single clone for each biological replicate (Methods), with dots representing experimental data.

**Extended Data Fig. 6.**
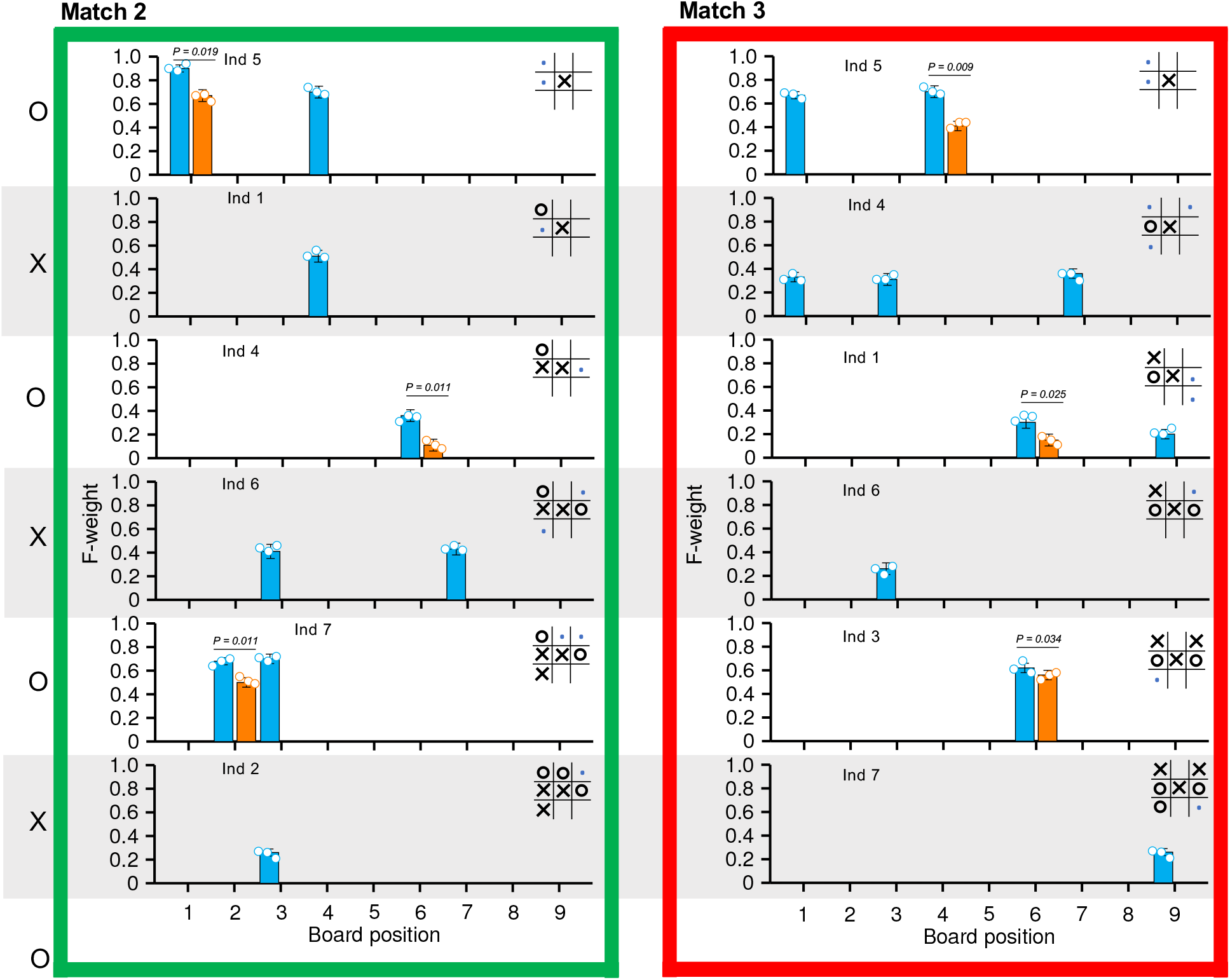
Detail of the remaining adaptation from Fig. 4. Grey overlays align starting with the second overlay of Fig. 4e.

**Extended Data Fig. 7.**
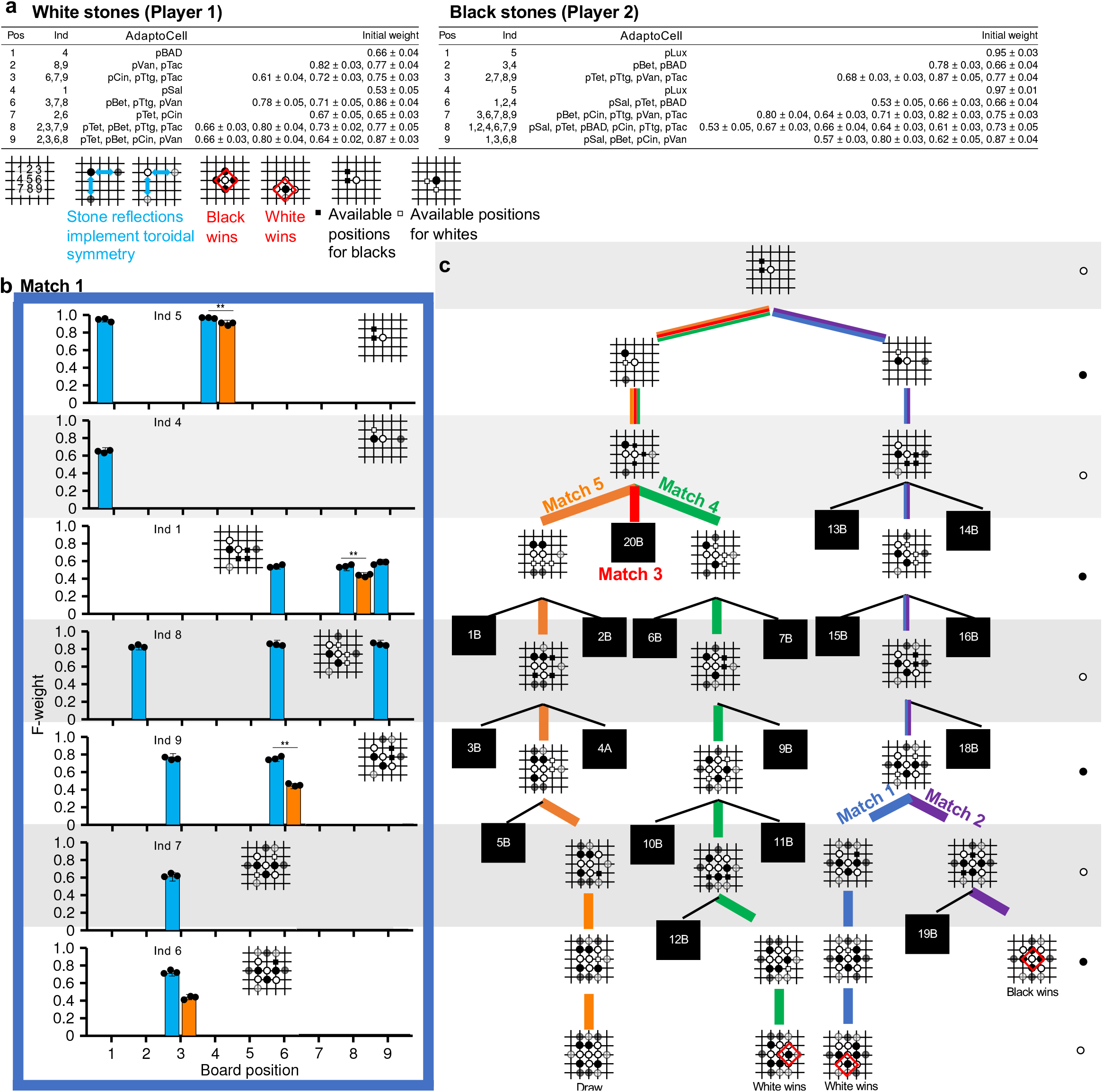
Bacteria cultures adapt playing and Adaptation Toroidal Atari Go against one other. **a**. AdaptoCell library used in the bacterial cultures at each position (pos) with several AdaptoCell types per position. Bacteria play and adapt against each other according to the decision tree in panel **c**. Weights were measured before adaptation as F-weights and by DNA sequencing. Stone reflections implement the toroidal symmetry by replicating the stones to all equivalent positions (positions differing by 3 rows or columns). A player triumphs by ’surrounding’ an opponent’s stone with their stones or replicas. **b,** Loss of the bacteria first player (white stones) leads to adaptation with kanamycin (normalised mCherry fluorescence fraction before (blue) and after (orange) negative reinforcement Adaptation (± s.d. for n = 3)). **c,** Game decision tree, with black boxes indicating collapsed branches. Coloured branches indicate the losses used as training steps for either player.

**Extended Data Fig. 8.**
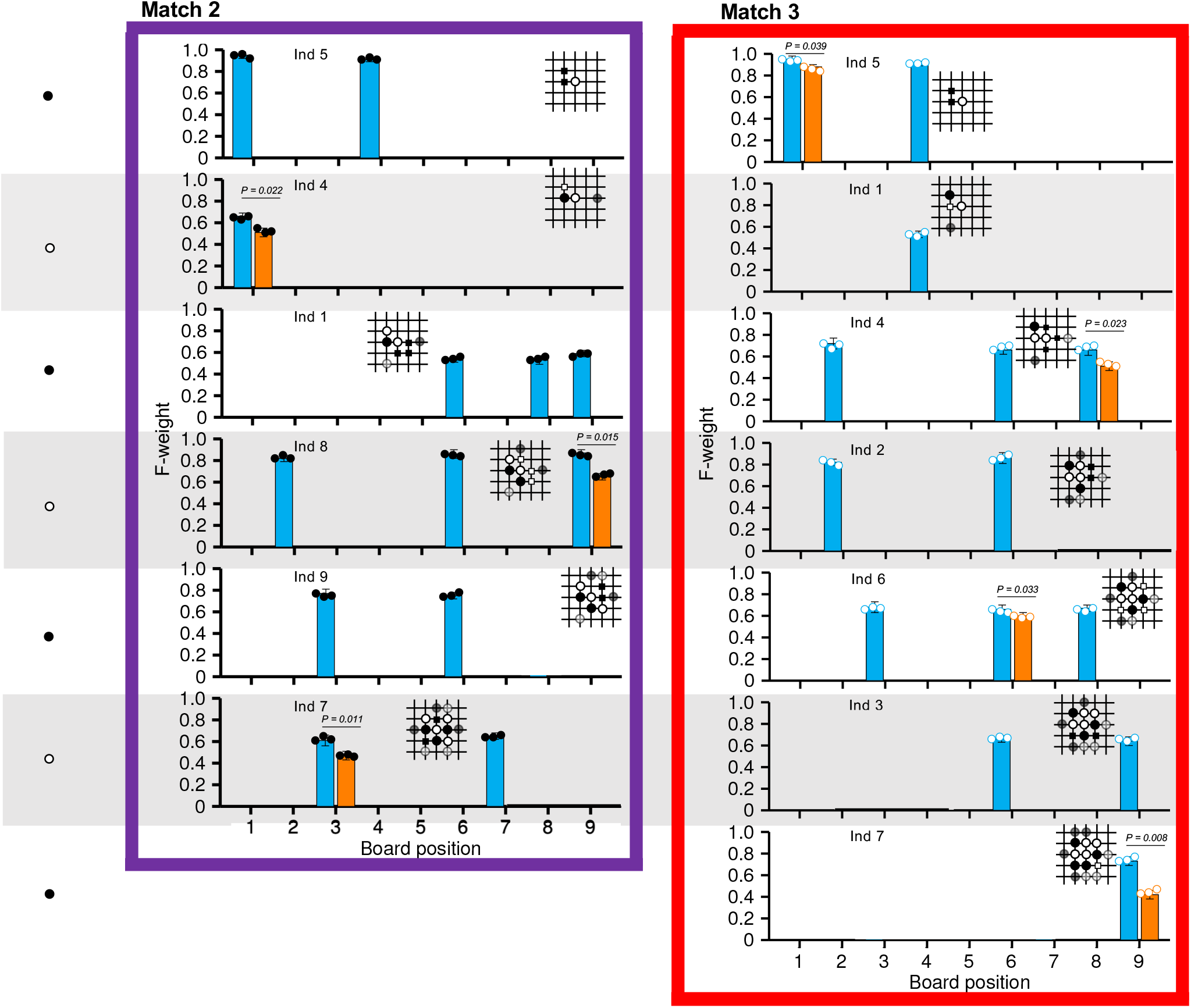
**Extended Data Fig. 7 continued.** Matches 2 and 3 of the tournament of Extended Data Fig. 7. Grey overlays align with the second overlay of Extended Data Fig. 7c.

**Extended Data Fig. 9.**
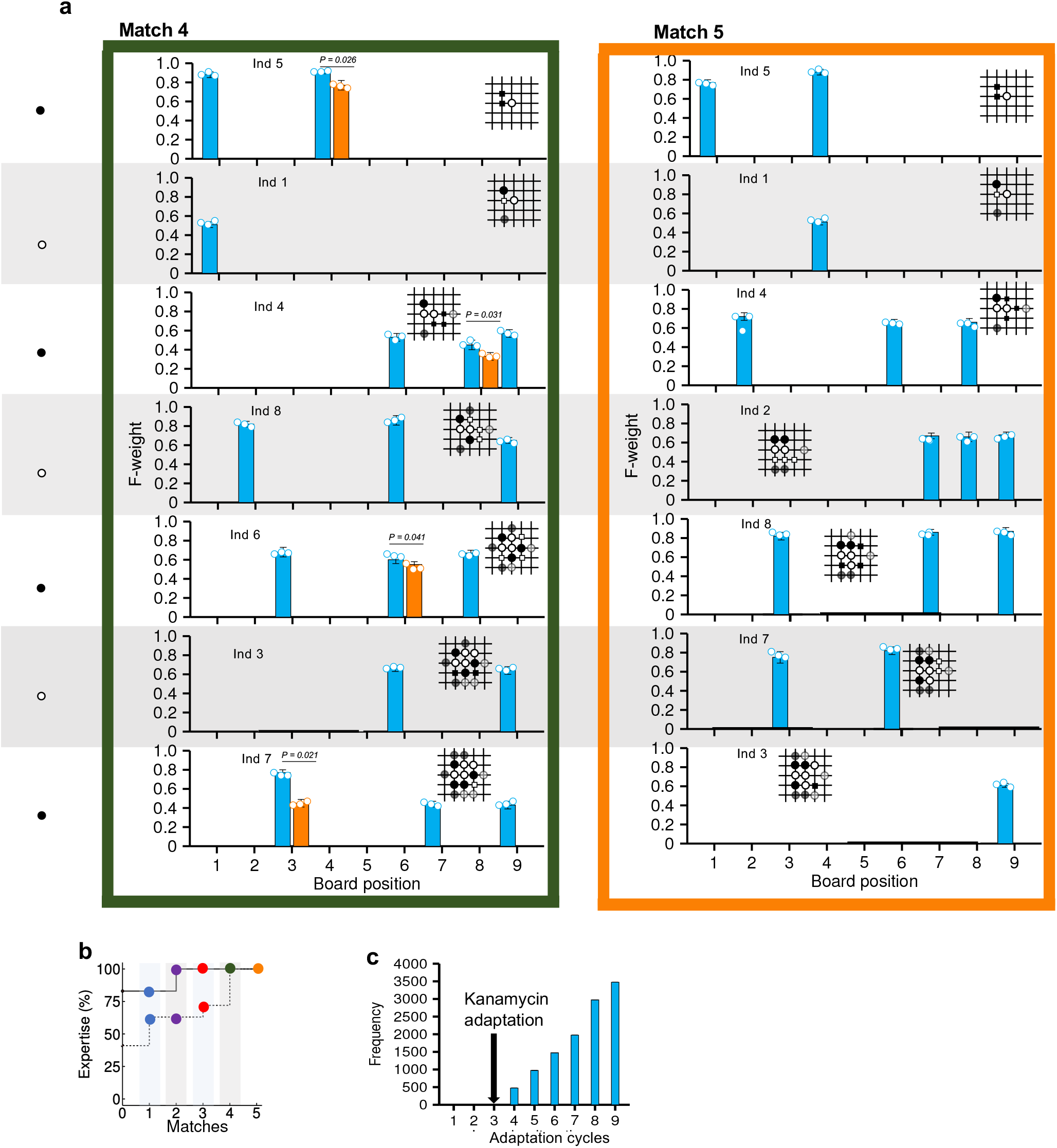
**Extended Data Fig. 7 continued. a,** Matches 5 and 6 of the tournament of Extended Data Fig. 7. Grey overlays align with the second overlay of Extended Data Fig. 7c. **b,** Evolution of the expertise during a tournament where both players are able to learn by kanamycin adaptation, with dotted line denoting player 2. Grey overlays indicate the moves of Player 1. **c,** Distribution of Adaptation rounds required to achieve mastery of a set of 10,000 simulations of tournaments where the adaptation uses randomised inducers (similarly to Fig. 1g).

**Extended Data Fig. 10.**
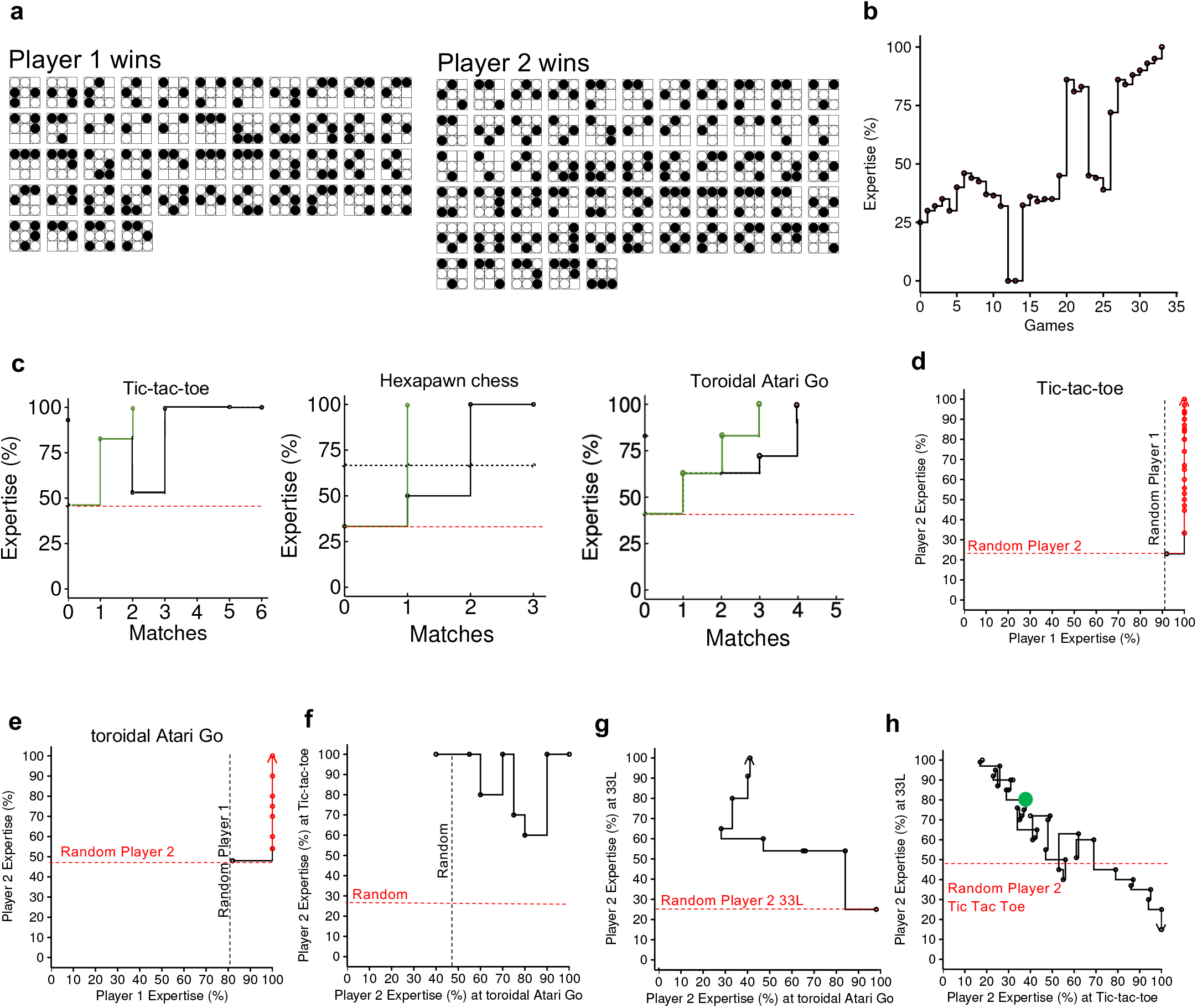
Testing of the capabilities of a 9YES library to re-learn new games by computer simulations. **a,** Rules of the 33L game (the most difficult to learn). **b,** Evolution of the expertise of Player 2 during a tournament where both players are able to learn the 33L game by kanamycin adaptation. Dots indicate the expertise after kanamycin adaptation. **c,** Comparison of our reinforcement Adaptation (black) of Fig. 3, 4 and Ext. Data Fig. 7-9 against the fastest possible Adaptation (green) for the 3 games studied experimentally in the paper. **d,** Expertise evolution as both Player 1 (black) and Player 2 (red) at the game of Tic-tac-toe. **e,** Expertise evolution as both Player 1 (black) and Player 2 (red) at the game of Toroidal Atari Go after Adaptation the game of Tic-tac-toe. The expertise of Tic-tac-toe was maintained while achieving mastery of Atari Go. **g,** Expertise evolution as Player 2 at the 33L game at the expense of losing the expertise at Atari Go (and Tic-tac-toe). **h,** Re-training of the library to achieve back its mastery as Player 2 at the game of Tic-tac-toe. Green marker indicates the AdaptoCell fusion at match 9. See more details in Supplementary Text.

